# Experience-dependent plasticity in an innate social behavior is mediated by hypothalamic LTP

**DOI:** 10.1101/2020.07.21.214619

**Authors:** Stefanos Stagkourakis, Giada Spigolon, Grace Liu, David J. Anderson

**Affiliations:** Division of Biology and Biological Engineering 156-29, Tianqiao and Chrissy Chen Institute for Neuroscience, California Institute of Technology, Pasadena, California 91125, USA; Biological Imaging Facility, California Institute of Technology, Pasadena, California 91125, USA; Howard Hughes Medical Institute, California Institute of Technology, 1200 East California Blvd, Pasadena, California 91125, USA

**Keywords:** innate behaviors, long term potentiation, ventromedial hypothalamus, testosterone

## Abstract

All animals can perform certain survival behaviors without prior experience, suggesting a “hard wiring” of underlying neural circuits. Experience, however, can alter the expression of innate behaviors. Where in the brain and how such plasticity occurs remains largely unknown. Previous studies have established the phenomenon of “aggression training,” in which the repeated experience of winning successive aggressive encounters across multiple days leads to increased aggressiveness. Here we show that this procedure also leads to long-term potentiation (LTP) at an excitatory synapse, derived from the Anterior Hippocampus/Posterior Medial amygdala (AHiPM), onto estrogen receptor 1-expressing (Esr1^+^) neurons in the ventrolateral subdivision of the ventromedial hypothalamus (VMHvl). We demonstrate further that the optogenetic induction of such LTP *in vivo* facilitates, while optogenetic long-term depression (LTD) diminishes, the behavioral effect of aggression training, implying a causal role for potentiation at AHiPM➔VMHvl^Esr1^ synapses in mediating the effect of this training. Interestingly, ∼25% of inbred C57BL/6 mice fail to respond to aggression training. We show that these individual differences are correlated both with lower levels of testosterone, relative to mice that respond to such training, and with a failure to exhibit LTP *in vivo* after aggression training. Administration of exogenous testosterone to such non-aggressive mice restores both behavioral and physiological plasticity *in vivo*. Together, these findings reveal that LTP at a hypothalamic circuit node mediates a form of experience-dependent plasticity in an innate social behavior, and a potential hormone-dependent basis for individual differences in such plasticity among genetically identical mice.

**Significance Statement:** Modification of instinctive behaviors occurs through experience, yet the mechanisms through which this happens have remained largely unknown. Recent studies have shown that potentiation of aggression, an innate behavior, can occur through repeated winning of aggressive encounters. Here we show that synaptic plasticity at a specific excitatory input to a hypothalamic cell population is correlated with, and required for, the expression of increasingly higher levels of aggressive behavior following successful aggressive experience. We additionally show that the amplitude and persistence of long-term potentiation at this synapse are influenced by serum testosterone, administration of which can normalize individual differences among genetically identical inbred mice, in the expression of intermale aggression.

## Introduction

Brains evolved to optimize the survival of animal species by generating appropriate behavioral responses to both stable and unpredictable features of the environments in which they live. Accordingly, two major brain strategies for behavioral control have been selected. In the first, developmentally specified neural circuits generate rapid innate responses to sensory stimuli that have remained relatively constant and predictable over evolutionary timescales (1-5). In the second, neural circuits generate flexible responses to stimuli that can change over an individual’s lifespan, through learning and memory (6-8).

One prevailing view is that these two strategies are implemented largely through distinct neuroanatomical structures and neurophysiological mechanisms. According to this view, in the mammalian brain innate behaviors are mediated by evolutionarily ancient, deep subcortical structures, such as the medial amygdala and hypothalamus, which link specific sensory inputs to evolutionarily “prepared” motor outputs through relatively stable synaptic connections (9-13). In contrast, learned behaviors are mediated by more recently evolved structures, such as the cortex and hippocampus, which compute flexible input-output mapping responses through synaptic plasticity mechanisms (14-19).

The idea that innate vs. learned behaviors are mediated by largely distinct neural systems has been reinforced by studies that have revealed, for example, distinct anatomical pathways through which olfactory cues evoke learned vs. innate behaviors in both the mouse (20-22) and in *Drosophila* (23-25); reviewed in (26). The concept of a dichotomous nervous system architecture for mediating appropriate biological responses to evolutionarily ancient vs. novel stimuli is analogous to the innate vs. adaptive branches of the immune system (27).

This view of distinct neural pathways for innate vs. learned behaviors, however, is challenged by case of behaviors that, while apparently “instinctive”, can nevertheless be modified by experience. For example, while aggression has been considered by classical ethologists as a prototypical “released” innate behavior (28, 29), studies from the 1940’s onwards showed that mice could be trained to be more aggressive by repeated fighting experience (30-35). Similarly, studies in rodents have shown that defensive behaviors such as freezing can be elicited by both unconditional and conditional stimuli, the latter via Pavlovian associative learning (reviewed in (36).

These observations raise the question of where in the brain, and how, the neural circuitry that mediates innate behaviors is modified by experience. In the case of defensive behaviors, such as freezing, the prevailing view argues for parallel pathways: conditioned defensive behavior is mediated by circuitry involving the hippocampus, thalamus and the basolateral/central amygdala, whereas innate defensive responses to predators are mediate by the medial amygdala (MeA)/bed nucleus of the stria terminalis (BNST) and hypothalamic structures (reviewed in (37)). Although the basolateral amygdala contains representations of unconditioned aversive and appetitive stimuli, these representations are used as the cellular substrate for pairing with conditioned stimuli (38, 39). Despite this segregation of learned and innate defensive pathways, it remains possible that experience-dependent influences on other innate behaviors may involve plasticity at synapses that directly mediate such behaviors (e.g., at inputs to midbrain PAGvl neurons (40)).

We have investigated this issue using inter-male offensive aggression in mice as an experimental paradigm. While mice housed under appropriate conditions can exhibit aggression in the absence of any prior agonistic encounters with male conspecifics (31, 33, 41), an effect of repeated successful aggressive experience to facilitate, or “prime”, subsequent attack behavior, considered as a form of “aggression training”, has been well-documented (42, 43). The neural substrate and physiological mechanisms underlying this form of experience-dependent plasticity of an innate behavior remain unknown. Interestingly, inbred strains of laboratory mice exhibit individual differences in the ability to manifest this form of behavioral plasticity, with up to 25% of animals failing to respond to aggression training (35). The biological basis of this apparent epigenetic heterogeneity is not understood. Here we provide data supporting a plausible explanation for both observations, one that links physiological plasticity at hypothalamic synapses to aggressive behavior and sex hormone levels.

## Results

### Aggression training increases VMHvl^Esr1^ neuron activity

Aggression levels escalate following the recurrent manifestation of the behavior (35, 44, 45), an effect termed here as “aggression training”. Using a five consecutive-day resident-intruder assay (5cdRI, Fig. 1A), aggression training was investigated in a cohort of C57BL/6 Esr1-Cre mice (n=138), which displayed increased aggression levels that remained significantly elevated, relative to pre-training animals, over a prolonged period of time (maximal period tested – three months, Fig. 1C-F). This assay enabled the identification of socially naive, aggressive (AGG), and non-aggressive (NON) mice, the latter of which represent ∼23% of all males tested (Fig. 1B). Aggression levels were found to plateau on the fourth and fifth day of the 5cdRI, suggestive of a ceiling effect in the expression of aggressive behavior (Fig. 1C-E). Interestingly, aggression levels remained elevated for the maximal follow-up period tested (three months) following aggression training, as compared to the first instance of resident-intruder (RI) test (Fig. 1C).

**Fig. 1.**
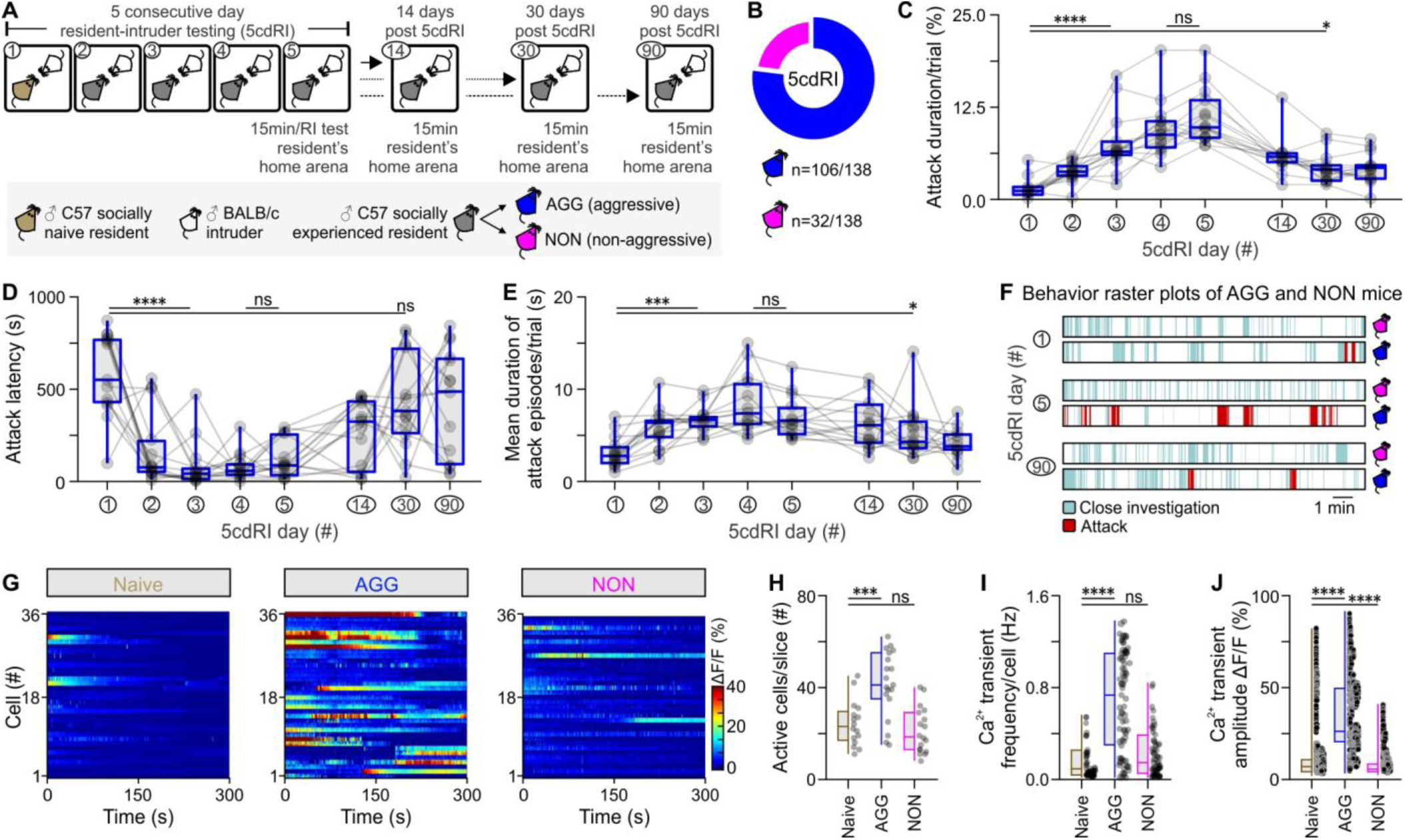
Aggression learning alters baseline activity dynamics in VMHvl^Esr1^ neurons. (A) Schematic of the experimental design - five consecutive day resident-intruder test (5cdRI), with three follow-up dates, used for the study of the behavioral effect of aggression training. (B) Summary indicative of the number of male animals exhibiting the two distinct aggression phenotypes (n=138). (C) Quantification of the cumulative duration (in %) of aggression per trial (n=15 AGG mice per group, Kruskal-Wallis one-way ANOVA with uncorrected Dunn’s post hoc test, *P* < 0.0001 between day 1 and day 3 of the 5cdRI, *P* = 0.4819 between day 4 and day 5 of the 5cdRI, *P* = 0.0149 between day 1 and day 30). (D) Quantification of attack latency (in seconds) of aggression per trial (n=15 AGG mice per group, Kruskal-Wallis one-way ANOVA with uncorrected Dunn’s post hoc test, *P* < 0.0001 between day 1 and day 3 of the 5cdRI, *P* = 0.5602 between day 4 and day 5 of the 5cdRI, *P* = 0.2184 between day 1 and day 30). (E) Quantification of average attack episode duration (in seconds) per trial (n=15 AGG mice per group, Kruskal-Wallis one-way ANOVA with uncorrected Dunn’s post hoc test, *P* < 0.0001 between day 1 and day 3 of the 5cdRI, *P* = 0.3326 between day 4 and day 5 of the 5cdRI, *P* = 0.0209 between day 1 and day 30). (F) Behavior raster plots from AGG and NON mice, at different days of the 5cdRI test. (G) Baseline Ca^2+^ activity of VMHvl^Esr1^ neurons recorded *ex vivo*, in brain slices of socially naive, aggressive (AGG), and non-aggressive (NON) males. (H) Quantification of active cells/slice (n=16-19 brain slices, collected from n=7-9 mice, one-way ANOVA with Dunnett’s post hoc test, *P* = 0.0002 between socially naive and AGG mouse brain slices, *P* = 0.6358 between socially naive and NON mouse brain slices). (I) Quantification of Ca^2+^ spike frequency per cell (n=16-19 brain slices, collected from n=7-9 mice, Kruskal-Wallis one-way ANOVA with Dunn’s post hoc test, *P* < 0.0001 between socially naive and AGG mice, *P* = 0.3331 between socially naive and NON mice). (J) Quantification of Ca^2+^ spike amplitude (n=16-19 brain slices, collected from n=7-9 mice, Kruskal-Wallis one-way ANOVA with Dunn’s post hoc test, *P* < 0.0001 between socially naive and AGG mice, *P* < 0.0001 between socially naive and NON mice). ns; not significant, \**P* < 0.05, \*\*\**P* < 0.001, \*\*\*\**P* < 0.0001. In box-and-whisker plots, center lines indicate medians, box edges represent the interquartile range, and whiskers extend to the minimal and maximal values.

To test whether aggression training involves plasticity in a structure that mediates the innate aspect of aggression (46), we initially focused on VMHvl^Esr1^ neurons, optogenetic stimulation of which can evoke attack in socially naïve, inexperienced animals (47). Using brain slice Ca^2+^ imaging, the average baseline activity of VMHvl^Esr1^ neurons was found to increase in AGGs but not in NONs, following aggression training (Fig. 1G-J). Voltage-clamp *ex vivo* VMHvl^Esr1^ neuron recordings revealed in AGG mice a significant increase in the frequency and amplitude of spontaneous excitatory postsynaptic currents (sEPSCs, Fig. 2A-C), relative to socially naïve animals. The increase in the amplitude of sEPSCs raised the possibility that a synaptic potentiation mechanism may be present in VMHvl^Esr1^ neurons (48, 49). In contrast, voltage-clamp *ex vivo* VMHvl^Esr1^ neuron recordings in slices from NON mice revealed an increase in the frequency and amplitude of spontaneous inhibitory postsynaptic currents (sIPSCs, Fig. S1).

**Fig. 2.**
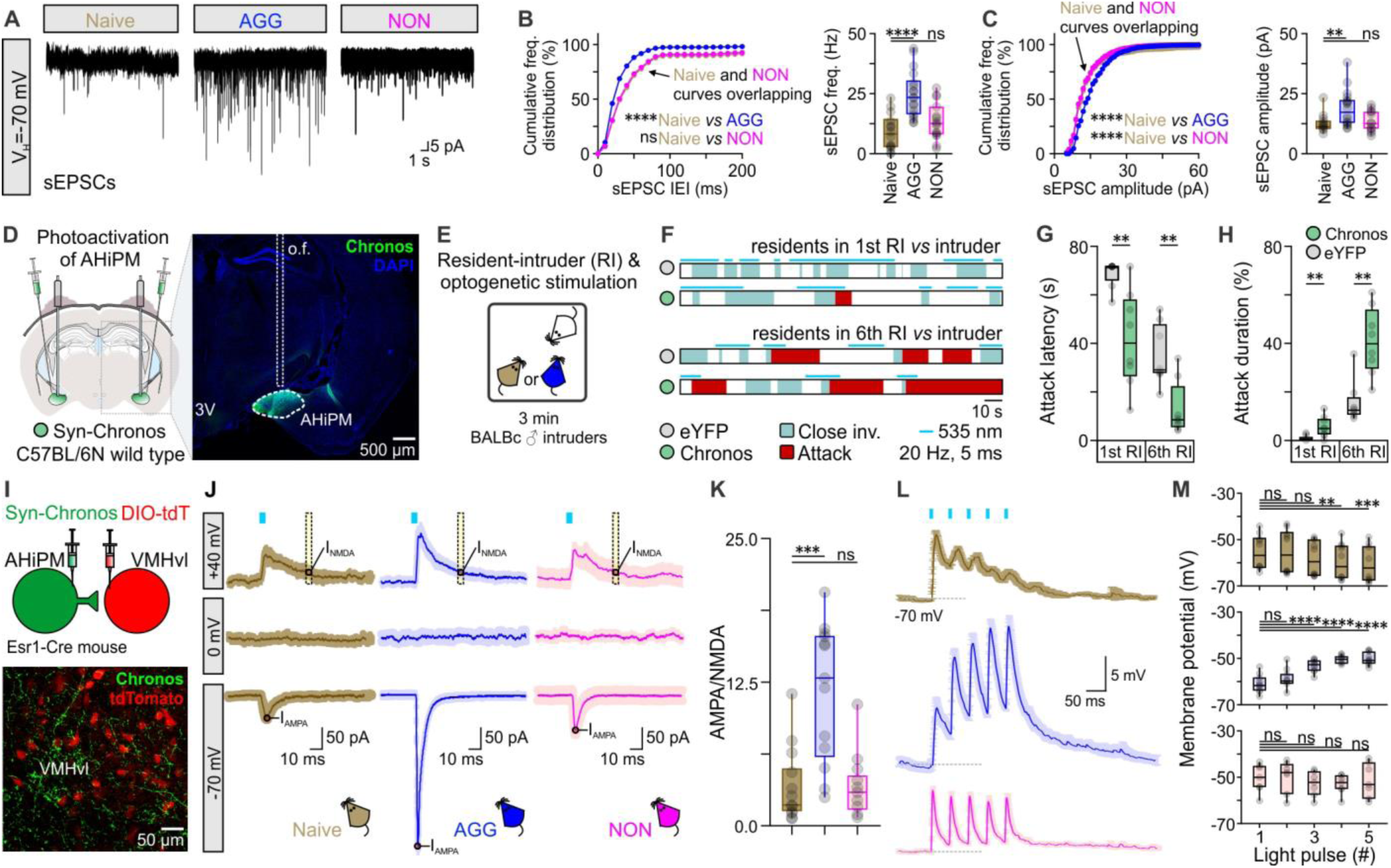
AHiPM➔VMHvl^Esr1^ synapses become potentiated following aggression training. (A) Representative recordings of spontaneous excitatory post-synaptic currents (sEPSCs) from VMHvl^Esr1^ neurons, from socially naive, AGG and NON mice. (B) Left – cumulative frequency distribution plot of sEPSC IEI in voltage-clamp recordings collected from VMHvl^Esr1^ neurons from socially naive, AGG and NON mice (n=14-18 VMHvl^Esr1^ neuron recording per group, collected from 8-10 mice per group, Kolmogorov-Smirnov test, *P* < 0.0001 between socially naive and AGG mice, *P* = 0.3454 between socially naive and NON mice). Right – comparison of sEPSC frequency from voltage-clamp recordings collected from VMHvl^Esr1^ neurons from socially naive, AGG and NON mice (n=14-18 VMHvl^Esr1^ neuron recording per group, collected from 8-10 mice per group, one-way ANOVA with Dunnett’s post hoc test, *P* < 0.0001 between socially naive and AGG mouse brain slices, *P* = 0.2576 between socially naive and NON mouse brain slices). (C) Left – cumulative frequency distribution plot of sEPSC amplitude in voltage-clamp recordings collected from VMHvl^Esr1^ neurons from socially naive, AGG and NON mice (n=14-18 VMHvl^Esr1^ neuron recording per group, collected from 8-10 mice per group, Kolmogorov-Smirnov test, *P* < 0.0001 between socially naive and AGG mice, *P* < 0.0001 between socially naive and NON mice). Right – comparison of sEPSC frequency from voltage-clamp recordings collected from VMHvl^Esr1^ neurons from socially naive, AGG and NON mice (n=14-18 VMHvl^Esr1^ neuron recording per group, collected from 8-10 mice per group, Kruskal-Wallis one-way ANOVA with uncorrected Dunn’s post hoc test, *P* = 0.0041 between socially naive and AGG mouse brain slices, *P* = 0.6712 between socially naive and NON mouse brain slices). (D) Left – schematic of the experimental design used for optogenetic studies of aggression following photoactivation of AHiPM, and right – confocal image indicative of Chronos-eYFP expression in AHiPM. (E) Schematic illustration of the experimental protocol used in AHiPM^Chronos^ stimulation experiments. (F) Sample behavior raster plots with *in vivo* optogenetics and social behavior in the resident intruder (RI) assay, of socially naive and AGG mice. (G) Quantification of attack latency, in the first and sixth RI trial (n=8 mice per group, first RI, two-sided Mann–Whitney U test, *P* = 0.0033 between YFP and Chronos groups, sixth RI, two-tailed unpaired *t*-test, *P* = 0.0022 between YFP and Chronos groups). (H) Quantification of attack duration, in the first and sixth RI trial (n=8 mice per group, first RI, two-sided Mann–Whitney U test, *P* = 0.0079 between YFP and Chronos groups, sixth RI, *P* = 0.0011 between YFP and Chronos groups). (I) Top – schematic of the experimental design used for the study of the AHiPM VMHvl synapse and bottom – confocal image indicative of AHiPM originating processes in VMHvl. (J) Identification of the AHiPM➔VMHvl synapse as purely excitatory, and extraction of the AMPA to NMDA ratio in socially naive, AGG and NON mice (average of n=13-14 neuron recordings from 8-9 socially naive, AGG and NON mice respectively). (K) Quantification of the AMPA/NMDA ratio (n=13-14, Kruskal-Wallis one-way ANOVA with Dunn’s post hoc test, *P* = 0.0005 between socially naive and AGG mice, *P* > 0.9999 between socially naive and NON mice). (L) Synaptic integration in VMHvl^Esr1^ neurons from socially naive, AGG and NON mice (average traces of n=7-10 neuron recordings from 7-9 mice respectively). (M) Quantification of the five optically-evoked excitatory post-synaptic potentials (oEPSP) peak amplitude presented in (O). Top - oEPSP amplitude quantification in VMHvl^Esr1^ neurons recorded from socially naive mice (n=10 neurons from 9 mice, Friedman one-way ANOVA with Dunn’s post hoc test, *P* > 0.9999 between 1^st^ and 2^nd^ pulse, *P* = 0.3587 between 1^st^ and 3^rd^ pulse, *P* = 0.0028 between 1^st^ and 4^th^ pulse, and *P* = 0.0009 between 1^st^ and 5th pulse). Middle - oEPSP amplitude quantification in VMHvl^Esr1^ neurons recorded from AGG mice (n=10 neurons from 9 mice, one-way ANOVA with Dunnett’s post hoc test, *P* = 0.2935 between 1^st^ and 2^nd^ pulse, *P* < 0.0001 between 1^st^ and 3^rd^ pulse, *P* < 0.0001 between 1^st^ and 4^th^ pulse, and *P* < 0.0001 between 1^st^ and 5th pulse). Bottom - oEPSP amplitude quantification in VMHvl^Esr1^ neurons recorded from NON mice (n=7 neurons from 7 mice, one-way ANOVA with Dunnett’s post hoc test, *P* = 0.9865 between 1^st^ and 2^nd^ pulse, *P* = 0.5704 between 1^st^ and 3^rd^ pulse, *P* = 0.0751 between 1^st^ and 4^th^ pulse, and *P* = 0.9803 between 1^st^ and 5th pulse). ns; not significant, \*\**P* < 0.01, \*\*\**P* < 0.001, \*\*\*\**P* < 0.0001. In box-and-whisker plots, center lines indicate medians, box edges represent the interquartile range, and whiskers extend to the minimal and maximal values.

The increased activity of VMHvl^Esr1^ neurons following aggression training in AGG mice prompted us to investigate whether this might involve potentiation of an excitatory input to these cells. Anatomical studies have identified a strong purely excitatory input to VMHvl^Esr1^ cells, which originates anatomically from the posteromedial part of the amygdalohippocampal area (AHiPM, also termed the posterior amygdala, PA) (50). *In vivo* optogenetic activation of AHiPM evoked aggression (Fig. 2D-H), confirming a recent report using chemogenetic activation (51). We found, moreover, that this effect is amplified in aggression-experienced animals (Fig. 2D-H). Investigation of the functional connectivity at AHiPM➔VMHvl^Esr1^ synapses in acute slices using optogenetic activation of AHiPM inputs (Fig. 2I-K) indicated that this projection is entirely excitatory, at least in part monosynaptic (Fig. S2), with an absence of any evoked responses at the reversal potential for excitation (Vm hold = 0 mV, Fig. 2J – middle row) and reliable photostimulation-evoked currents at the reversal potential for inhibition (Vm hold = −70 mV, Fig. 2J – bottom row). These observations raised the question of whether potentiation at AHiPM➔VMHvl^Esr1^ synapses underlies the observed increase in the excitatory synaptic input onto VMHvl^Esr1^ neurons recorded from AGG mice following aggression training.

### Plasticity at a hypothalamic synapse following aggression training

At the postsynaptic side of synapses that can undergo LTP, the response to stimulation of excitatory pre-synaptic inputs largely depends on the ratio of N-methyl-D-aspartate (NMDA) and α-amino-3-hydroxy-5-methyl-4-isoxazoleproprionic acid (AMPA) receptors (52, 53). To test whether the AHiPM➔VMHvl^Esr1^ synapse undergoes potentiation following aggression training, therefore, we measured the AMPA/NMDA ratio (54-56). This analysis revealed a significantly higher AMPA/NMDA ratio in AGGs following such training, compared to socially naive and NON mice (Fig. 2K). As the AMPA/NMDA ratio can influence synaptic integration properties (57), we also investigated synaptic integration in VMHvl^Esr1^ neurons from socially naive, AGG (trained) and NON mice. Indeed, these three groups of animals exhibited distinct synaptic integration properties, with depressing/static synaptic integration in socially naive and NON mice, and facilitating synaptic integration in the VMHvl^Esr1^ neurons of AGG (trained) mice (Fig. 2L, M).

Changes in the AMPA/NMDA ratio and synaptic integration properties are often accompanied by changes in neuronal morphology and dendritic spine complexity, which can be indicative of structural LTP (sLTP) (58-62). To investigate this possibility, VMHvl^Esr1^ neurons which exhibited an increase in the AMPA/NMDA ratio and synaptic integration in acute slice preparations were filled with neurobiotin, and super-resolution images for reconstruction were obtained using the Airyscanning technique in a ZEISS LSM 880 (63, 64). Analysis of second-order dendritic segments identified a prominent increase in dendritic arborizations in VMHvl^Esr1^ neurons from AGG (trained) mice, in comparison to socially naive and NON mice (Fig. 3A-L). These changes were reflected in most spine parameters measured, including density, branching points, volume, area, length, and mean diameter (Fig. 3M-R). However, the principal feature was an increase in the number of short-length spines, suggesting they were newly generated during or after training. Collectively, these observations suggested the possibility that potentiation is likely to occur at AHiPM➔VMHvl^Esr1^ synapses, following aggression training in susceptible animals. We therefore pursued this possibility using more specific electrophysiological protocols.

**Fig. 3.**
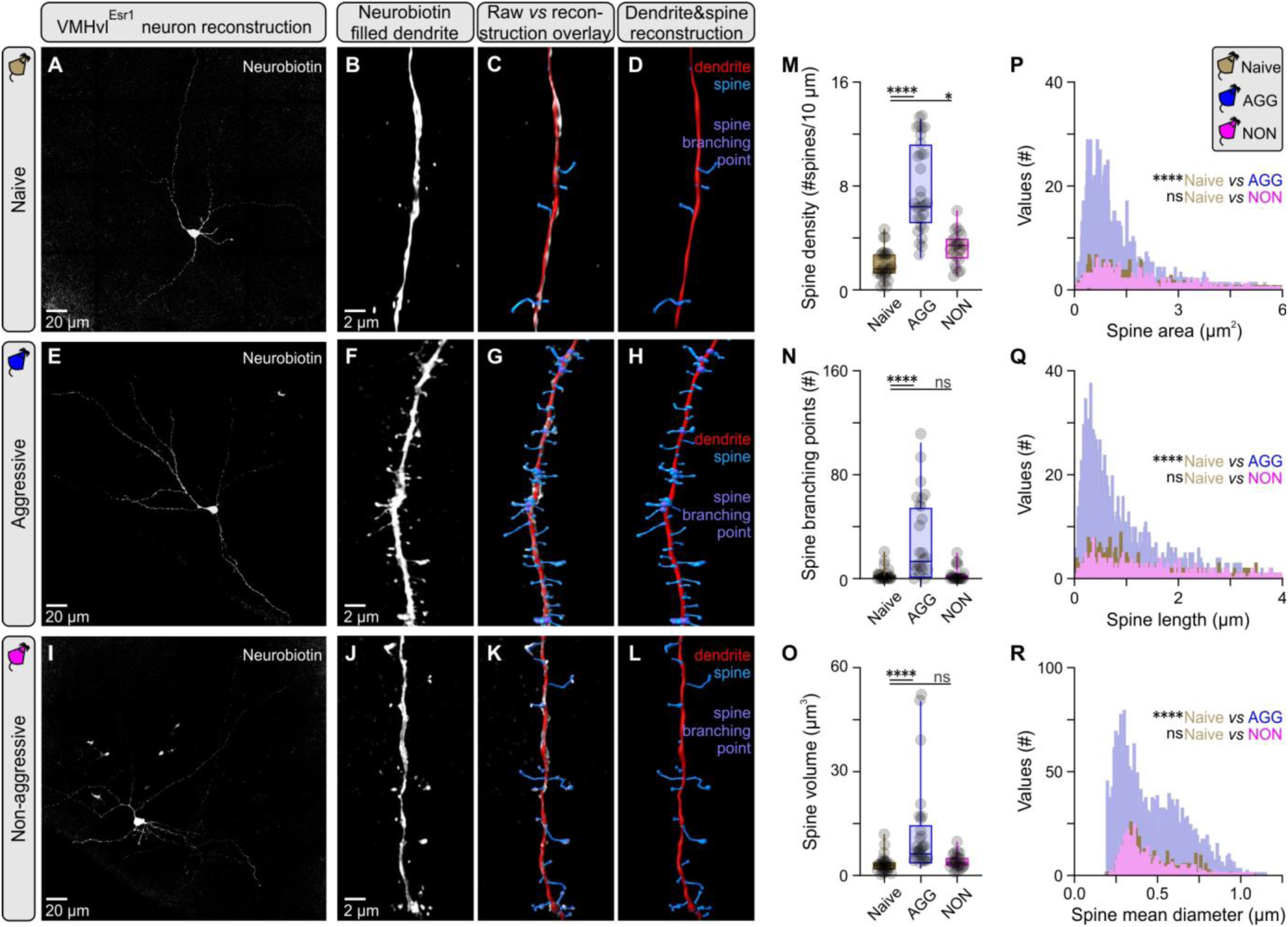
Increased dendritic spine complexity in VMHvl^Esr1^ neurons following aggression training. (A) Maximum projection confocal image of a VMHvl^Esr1^ neuron from a socially naive mouse recorded *ex vivo,* and filled with Neurobiotin. (B) 3D rendering of a second order dendritic segment Airyscan image from the neuron presented in (A). (C) Overlay of reconstruction data generated in Imaris against 3D rendering for the dendritic segment presented in (B). (D) Reconstructed dendritic segment of a VMHvl^Esr1^ neuron from a socially naive mouse, with color coding for the dendrite and spines. (E) Maximum projection confocal image of a VMHvl^Esr1^ neuron from an aggressive (AGG) mouse recorded *ex vivo,* and filled with Neurobiotin. (F) 3D rendering of a second order dendritic segment Airyscan image from the neuron presented in (E). (G) Overlay of reconstruction data generated in Imaris against 3D rendering for the dendritic segment presented in (F). (H) Reconstructed dendritic segment of a VMHvl^Esr1^ neuron from an AGG mouse, with color coding for the dendrite and spines. (I) Maximum projection confocal image of a VMHvl^Esr1^ neuron from a non-aggressive (NON) mouse recorded *ex vivo,* and filled with Neurobiotin. (J) 3D rendering of a second order dendritic segment Airyscan image from the neuron presented in (I). (K) Overlay of reconstruction data generated in Imaris against 3D rendering for the dendritic segment presented in (J). (L) Reconstructed dendritic segment of a VMHvl^Esr1^ neuron from a NON mouse, with color coding for the dendrite and spines. (M) Quantification of spine density in second order dendrites of VMHvl^Esr1^ neurons from socially naive, AGG and NON mice (n=3-5 cells per group, n=1 cell/brain slice/animal, n=23-26 segments analyzed per group, Kruskal-Wallis one-way ANOVA with Dunn’s post hoc test, *P* < 0.0001 between socially naive and AGG mice, *P* = 0.0432 between socially naive and NON mice). (N) Quantification of branching points in second order dendrites of VMHvl^Esr1^ neurons from socially naive, AGG and NON mice (n=23-26 segments analyzed per group, Kruskal-Wallis one-way ANOVA with Dunn’s post hoc test, *P* < 0.0001 between socially naive and AGG mice, *P* = 0.9969 between socially naive and NON mice). (O) Quantification of spine volume in second order dendrites of VMHvl^Esr1^ neurons from socially naive, AGG and NON mice (n=23-26 segments analyzed per group, Kruskal-Wallis one-way ANOVA with Dunn’s post hoc test, *P* < 0.0001 between socially naive and AGG mice, *P* = 0.4640 between socially naive and NON mice). (P) Frequency distribution plot of spine area, of spines present in second order dendrites in VMHvl^Esr1^ neurons from socially naive, AGG and NON mice (n=3-5 cells per group, n = 1 cell/brain slice/animal, n=402-2365 spines per group). (Q) Frequency distribution plot of spine length, of spines present in second order dendrites in VMHvl^Esr1^ neurons from socially naive, AGG and NON mice (n=402-2365 spines per group). (R) Frequency distribution plot of spine mean diameter, of spines present in second order dendrites in VMHvl^Esr1^ neurons from socially naive, AGG and NON mice (n=402-2365 spines per group). ns; not significant, \**P* < 0.05, \*\*\*\**P* < 0.0001. In box-and-whisker plots, center lines indicate medians, box edges represent the interquartile range, and whiskers extend to the minimal and maximal values.

### Experimental induction of LTP and LTD at AHiPM➔VMHvl^Esr1^ synapses

LTP and LTD can be experimentally induced in slices from brain areas typically associated with higher cognitive processing (65-70), such as the hippocampus, but few studies have demonstrated that this can occur in the hypothalamus (71-73), a site traditionally considered the source of instincts.

To determine whether LTP can be experimentally induced at AHiPM➔VMHvl^Esr1^ synapses *ex vivo,* we employed acute VMHvl slices, and used an optogenetics protocol composed of three bouts of photostimulation of Chronos-expressing AHiPM terminals, during which the VMHvl^Esr1^ neuron (identified by Cre-dependent expression of tdTomato in Esr1-Cre mice) was voltage-clamped at a depolarized membrane potential (−30 mV; Fig. 4A, 4B). The choice of this Hebbian stimulation protocol was based on our initial finding that combined pre- and post-synaptic depolarization were necessary for induction of potentiation at AHiPM➔VMHvl^Esr1^ synapses (Fig. S3) (74). The stimulation frequency of AHiPM terminals used here (20 Hz) was chosen based on our previous demonstration that direct optogenetic stimulation of VMHvl^Esr1^ neurons at this same frequency drives action potential firing with 100% spike fidelity *ex vivo* as well as *in vivo,* without depolarization block (47), and that it produces reliable synaptic integration in VMHvl^Esr1^ neurons (Fig. 2L, M). *Ex vivo* whole-cell voltage-clamp recordings composed of 20 min baseline and 20 min follow-up, revealed that the majority of VMHvl^Esr1^ neurons recorded in slices from socially naive, AGG (trained) and NON mice were able to express synaptic potentiation in response to this manipulation (Fig. 4C). Comparison of the responses with the animals’ aggression phenotypes revealed, however, that the dynamics of the response, including its maximum amplitude and persistence, differed between groups, with synaptic potentiation in slices from NON mice returning to baseline levels within the maximal period tested (20 min, Fig. 4D). Based on the Hebbian conditions required to evoke this form of synaptic potentiation, and the similarity of its features to LTP as characterized in the hippocampus (75, 76), we refer to this form of plasticity as hypothalamic LTP.

**Fig. 4.**
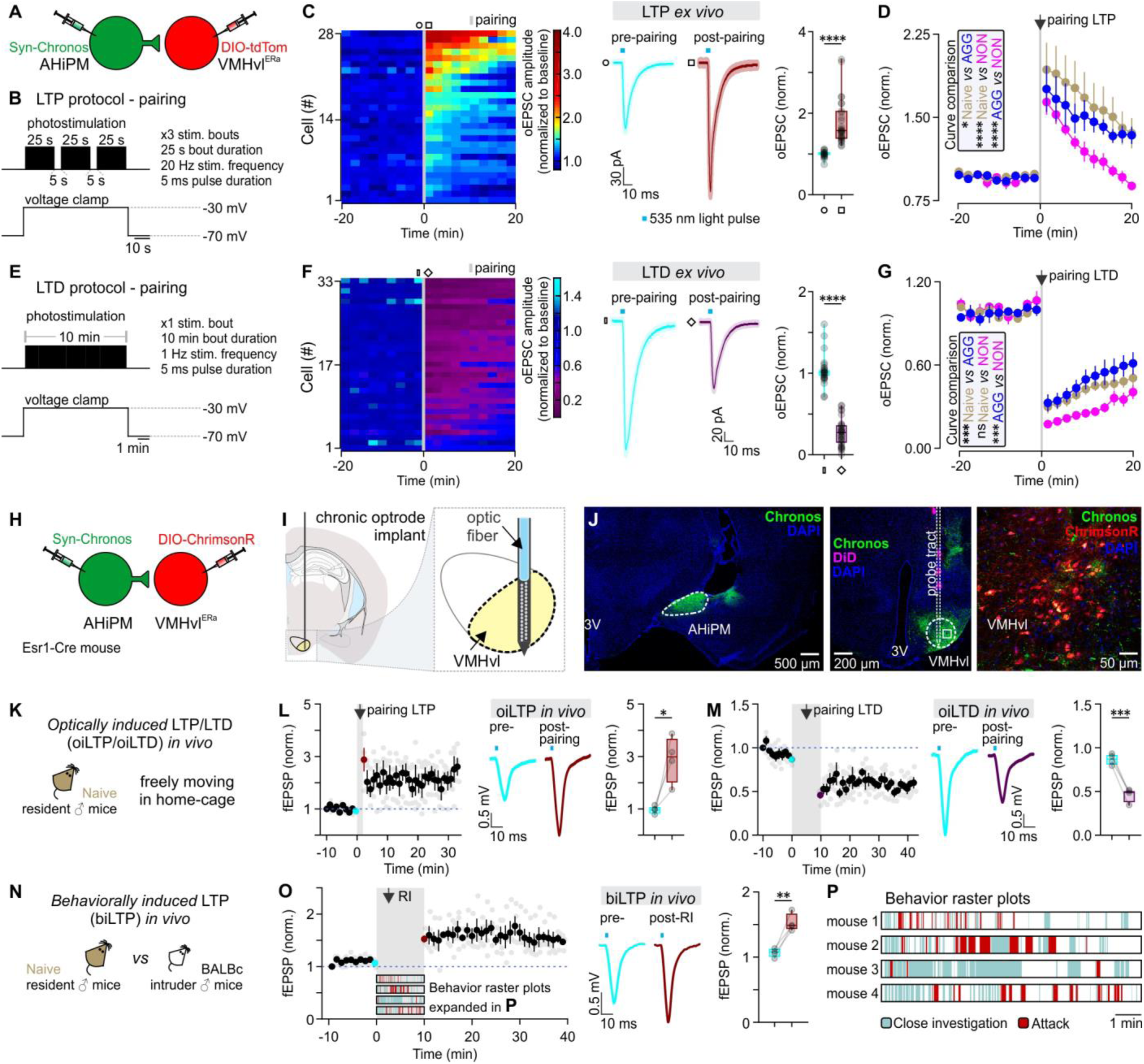
Induction of LTP and LTD at AHiPM➔VMHvl^Esr1^ synapses *ex vivo* and *in vivo*. (A) Schematic of the experimental design used to study the induction of LTP and LTD *ex vivo* in socially naive, aggressive (AGG) and non-aggressive (NON) mice. (B) Illustration of the experimental protocol used to induce LTP in the AHiPM➔VMHvl synapse. (C) Left – heat map illustrating the magnitude of LTP induction in all recorded VMHvl^Esr1^ neurons. Middle – average current immediately prior to and following the induction of LTP (middle, n=28 neurons collected from the three groups – socially naive, AGG and NON, with n=6-8 mice per group, light color envelope is the standard error). Right – quantification of the optically evoked excitatory post-synaptic current (oEPSC), prior to and following the induction of LTP (pre- *vs* post-pairing, n=28 neurons collected from the three groups – socially naive, AGG and NON, with n=6-8 mice per group, two-tailed Wilcoxon signed-rank test, *P* < 0.0001). (D) Identification of differences in amplitude and persistence of LTP in socially naive, AGG, and NON mice (n=9 neurons for 6 socially naive mice, n=10 neurons from 8 AGGs, and n=9 neurons from 6 NONs, Kolmogorov-Smirnov test for curve comparison, *P* = 0.0183 between socially naive and AGG mice, *P* < 0.0001 between socially naive and NON mice, and *P* < 0.0001 between AGG and NON mice). (E) Illustration of the experimental protocol used to induce LTD in the AHiPM➔VMHvl synapse. (F) Left – heat map illustrating the magnitude of LTD induction in all recorded VMHvl^Esr1^ neurons. Middle – average current immediately prior to and following the induction of LTD (middle, n=33 neurons collected from the three groups – socially naive, AGG and NON, with n=8 mice per group, light color envelope is the standard error). Right – quantification of the oEPSC, prior to and following the induction of LTD (pre- *vs* post-pairing, n=33 neurons collected from the three groups – socially naive, AGG and NON, with n=8 mice per group, two-tailed Wilcoxon signed-rank test, *P* < 0.0001). (G) LTD dynamics in the three groups (n=12 neurons for 8 socially naive mice, n=10 neurons from 8 AGGs, and n=11 neurons from 8 NONs, Kolmogorov-Smirnov test for curve comparison, *P* = 0.0002 between socially naive and AGG mice, *P* > 0.9999 between socially naive and NON mice, and *P* = 0.0008 between AGG and NON mice). (H) Schematic of the experimental design used to study the induction of LTP and LTD *in vivo* in socially naive mice. (I) Schematic illustration of the target coordinates of the optrode used to record local field potentials in VMHvl. (J) Left – representative confocal image of Chronos-eYFP expression in AHiPM. Middle - representative confocal image of the silicon probe tract targeted to VMHvl. Right – high magnification confocal image of VMHvl. (K) Illustration of the experimental design used to induce LTP or LTD in the AHiPM➔VMHvl synapse *in vivo*. (L) Left – plot of average of four experiments from four mice of field EPSP slope (normalized to baseline period) before and after optically-induced LTP (oiLTP). Middle – *in vivo* average field response prior to and following the induction of LTP. Right – quantification of optically induced field EPSPs (fEPSP), prior to and following the induction of LTP (pre- *vs* post-pairing, n=4 mice per group, two-tailed paired *t*-test, *P* = 0.0283). (M) Left – plot of average of four experiments from four mice of field EPSP slope (normalized to baseline period) before and after optically-induced LTD (oiLTD). Middle – *in vivo* average field response prior to and following the induction of LTD. Right – quantification of optically induced field EPSPs (fEPSP), prior to and following the induction of LTD (pre- *vs* post-pairing, n=4 mice per group, two-tailed paired *t*-test, *P* = 0.0007). (N) Illustration of the experimental design used to test the behavioral induction of LTP. (O) Left – plot of average of four experiments from four mice of field EPSP slope (normalized to baseline period) before and after behaviorally-induced LTP (biLTP). Middle – *in vivo* average field response prior to and following the behavioral induction of LTP. Right – quantification of optically induced field EPSPs (fEPSP), prior to and following social behavior experience in a socially naive mouse (pre- *vs* post-pairing, n=4 mice per group, two-tailed paired *t*-test, *P* = 0.0071). (P) Illustration of the behaviors expressed in the resident-intruder assay from socially naive mice used for the *in vivo* study of hypothalamic LTP. ns; not significant, \**P* < 0.05, \*\**P* < 0.01, \*\*\**P* < 0.001, \*\*\*\**P* < 0.0001. In box-and-whisker plots, center lines indicate medians, box edges represent the interquartile range, and whiskers extend to the minimal and maximal values.

These findings in turn raised the question of whether AHiPM➔VMHvl^Esr1^ synapses can also express long-term synaptic depression (LTD). This was investigated using a longer stimulation protocol for activating AHiPM terminals (10 min continuous stimulation at 1Hz, Fig. 4E). Similar to the case of LTP, most VMHvl^Esr1^ neurons expressed LTD of varying amplitude and dynamics, in a manner that varied with the animals’ aggression phenotypes (Fig. 4F, 4G). Interestingly, VMHvl^Esr1^ cells from NON mice expressed higher amplitude LTD (Fig. 4G) than did cells from other groups.

An important question raised by these *ex vivo* observations was whether LTP and LTD can be induced at AHiPM➔VMHvl^Esr1^ synapses *in vivo*, using either optogenetic stimulation or aggression training. To study the optogenetic induction of LTP *in vivo*, we used a similar paradigm to the *ex vivo* stimulation protocol (Fig. 4A, 4H). However, in order to be able to simultaneously depolarize both pre- and post-synaptic terminals we used spectrally segregated opsins with an overlap at 535 nm, to permit co-excitation (77). As in the case of the *ex vivo* experiments, AHiPM was transduced with Chronos; in addition, VMHvl^Esr1^ neurons were transduced with ChrimsonR (Fig. 4H). Chronic implantation of a silicon probe optrode in the VMHvl allowed the detection of optically induced LTP or LTD in individual freely-moving mice in their home cage, as a change in AHiPM stimulation-evoked local field potentials (fEPSPs; Fig. 4I-4K). Application of the Hebbian protocol in socially naive mice led to the robust expression of LTP in VMHvl *in vivo* (Fig. 4L), while application of the spaced protocol led to robust expression of LTD (Fig. 4M). Although VMHvl^Esr1^ neurons in silicon probe recordings were identified by optogenetic photo-tagging of post-synaptic cells, we cannot exclude that other classes of VMHvl neurons contribute to recorded fEPSPs.

These findings in turn raised the question of whether LTP can be induced *in vivo* by aggression training. Applying the same testing method used to analyze the optogenetic induction of *in vivo* LTP and LTD, the field excitatory postsynaptic potential (fEPSP) was monitored during a 10 min baseline period and then following aggression training in initially socially naive mice. We used test optogenetic pulses to briefly activate AHiPM terminals and ask whether the fEPSP increased in amplitude following the expression of aggression. Indeed, LTP was induced in VMHvl^Esr1^ neurons immediately after the expression of social behavior and aggression (Fig. 4N-P, n=4 mice tested). Notably, the behavioral induction of LTP (Fig. 4O), led to a persistent change in the amplitude of the fEPSP. This might suggest a lack of an early- *vs* late-phase distinction in the LTP at AHiPM➔VMHvl^Esr1^ synapses, in contrast to LTP features observed at defined synapses in the hippocampus and the amygdala (78, 79).

The above findings identify hypothalamic synaptic plasticity, and specifically LTP and LTD, as mechanisms that can occur and can alter VMHvl^Esr1^ neuronal excitability *in vivo*. Next we sought to address whether LTP and LTD have a causal role in the behavioral effect of aggression training.

### LTP facilitates and LTD inhibits potentiation of aggression following aggression training

To address whether LTP in the AHiPM➔VMHvl^Esr1^ synapses can influence the expression of aggression in inexperienced animals, we performed an *in vivo* optogenetic manipulation using the approach we established for the experimental induction of hypothalamic LTP *in vivo.* The AHiPM of Esr1-Cre mice was transduced with Chronos or YFP, while VMHvl^Esr1^ neurons were transduced with Cre-dependent ChrimsonR or mCherry (Fig. S4A). The effects of these manipulations were investigated behaviorally, not physiologically, therefore silicon recording probes were not implanted.

Following application of the LTP induction protocol over three consecutive days in socially naïve, solitary mice, the RI test was performed on the fourth day, and behavior was quantified (Fig. S4B-G). The opsin expressing mice (LTP group), exhibited elevated levels of aggression, indicating that the experimental induction of LTP in this particular projection *in vivo* can influence the expression of aggression in the absence of prior social experience (Fig. S4D-G).

The above paradigm was further modified to test, whether LTP and LTD can exert a causal influence on aggression training. Using the approach we established for the experimental induction of hypothalamic LTP or LTD *in vivo* (Fig. 4K-M), the AHiPM of Esr1-Cre mice was transduced with Chronos or YFP, while VMHvl^Esr1^ neurons were transduced with Cre-dependent ChrimsonR or mCherry (Fig. 5A) and the effects of these manipulations were investigated behaviorally.

**Fig. 5.**
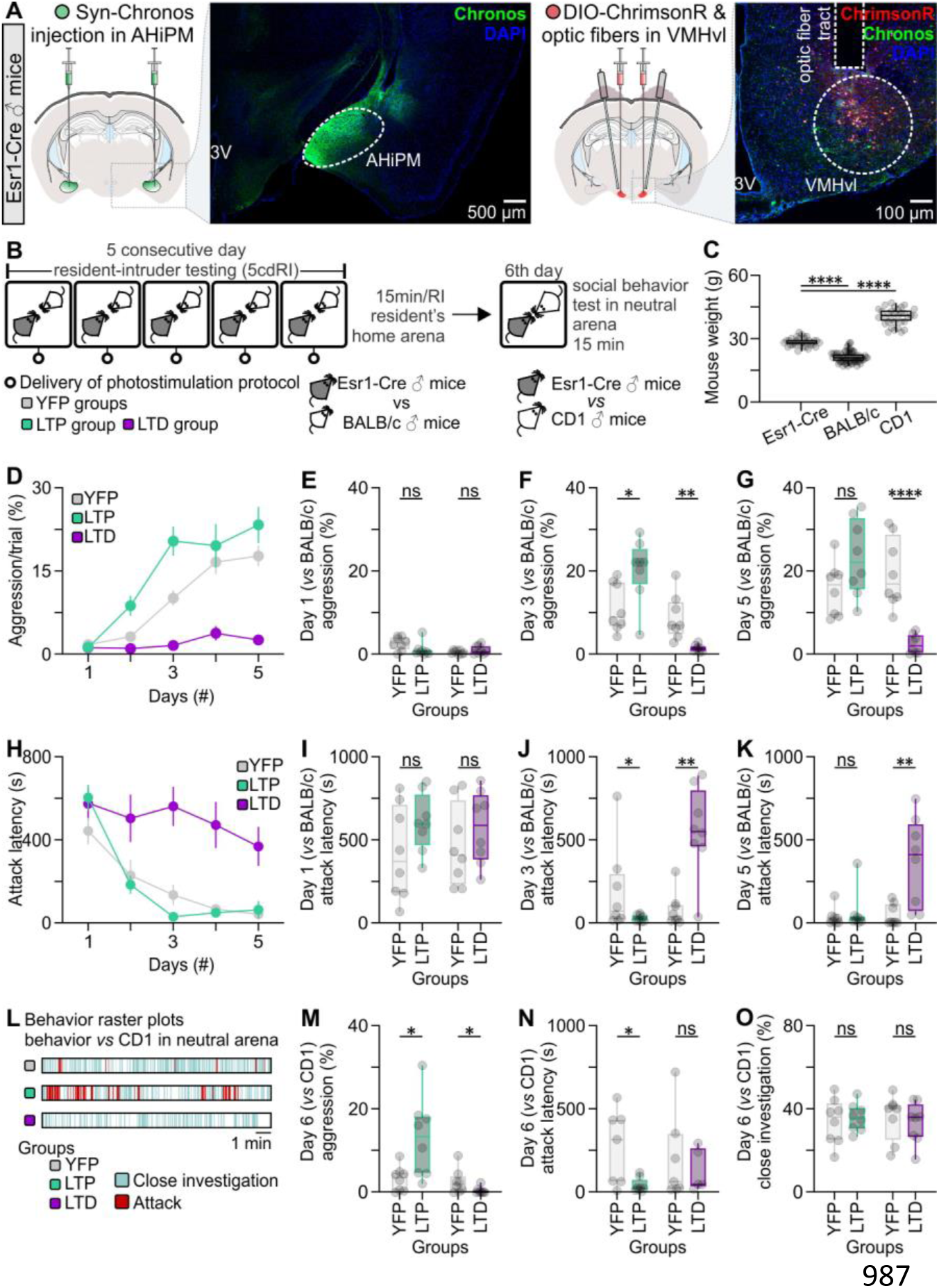
Optogenetic induction of LTP or LTD at AHiPM➔VMHvl^Esr1^ synapses *in vivo* facilitates or abolishes, respectively, the effect of aggression training. (A) Left-representative confocal image and schematic indicative of ChrimsonR expression in VMHvl^Esr1^ neurons, eYFP terminals of the AHiPM➔VMHvl projection, and the optic fiber tract terminating above VMHvl. Right - representative confocal image and schematic indicative of Chronos-eYFP expression in AHiPM. (B) Schematic of the experimental design used to identify whether LTP and LTD have an impact on aggression training. (C) Weight measurements of the mice which were used in the protocol; specifically, the Esr1-Cre mice were used as residents, the BALB/c as intruders, and the CD1 as novel conspecifics in a novel/neutral arena (n=32-64 mice per group, one-way ANOVA with Tukey’s test, *P* < 0.0001 between Esr1-Cre and BALB/c mice, *P* < 0.0001 between Esr1-Cre and CD1 mice). (D) Quantification of aggression levels expressed during a trial throughout the 5 consecutive day RI test (5cdRI) in the YFP (control), LTP and LTD groups. (E) Quantification of aggression levels on the first day of the 5cdRI test (n=8 mice per group, two-tailed unpaired *t*-test, *P* = 0.1049 between YFP and LTP groups, *P* = 0.2304 between YFP and LTD groups). (F) Quantification of aggression levels on the third day of the 5cdRI test (n=8 mice per group, two-tailed unpaired *t*-test, *P* = 0.0162 [observed power=0.989, Cohen’s D=0.7979, difference between means=9.13±3.34%, 95% CI =1.966 to 16.29] between YFP [lower 95% CI=6.452, higher 95% CI=16.06] and LTP [lower 95% CI=14.12, higher 95% CI=26.65] groups, *P* = 0.0017 between YFP and LTD groups). (G) Quantification of aggression levels on the fifth day of the 5cdRI test (n=8 mice per group, two-tailed unpaired *t*-test, *P* = 0.0777 between YFP and LTP groups, *P* < 0.0001 between YFP and LTD groups). (H) Quantification of attack latency throughout the 5cdRI in the YFP (control), LTP and LTD groups. (I) Quantification of attack latency on the first day of the 5cdRI test (n=8 mice per group, two-tailed unpaired *t*-test, *P* = 0.1406 between YFP and LTP groups, *P* = 0.3688 between YFP and LTD groups). (J) Quantification of attack latency on the third day of the 5cdRI test (n=8 mice per group, two-sided Mann–Whitney U test, *P* = 0.0415 [observed power=0.999, Cohen’s D=0.6072, difference between means=159.40±90.79 sec, 95% CI =-378.8 to 60.04] between YFP [lower 95% CI=-21.05, higher 95% CI=407.0] and LTP [lower 95% CI=16.54, higher 95% CI=50.56] groups, *P* = 0.0019 between YFP and LTD groups). (K) Quantification of attack latency on the fifth day of the 5cdRI test (n=8 mice per group, two-sided Mann–Whitney U test, *P* = 0.5054 between YFP and LTP groups, two-tailed unpaired *t*-test, *P* = 0.0052 between YFP and LTD groups). (L) Representative behavior raster plots of YFP, LTP and LTD mouse behavior in novel arena towards a novel CD1 conspecific. (M) Quantification of aggression levels on the sixth day against a CD1 male (n=8 mice per group, two-tailed unpaired *t*-test, *P* = 0.0387 [observed power=0.999, Cohen’s D=0.8980, difference between means=9.816±3.864%, 95% CI=0.6784 to 18.95] between YFP [lower 95% CI=0.8293, higher 95% CI=5.768] and LTP [lower 95% CI=5.190, higher 95% CI=21.04] groups, two-sided Mann–Whitney U test, *P* = 0.0295 [observed power=0.907, Cohen’s D=0.7357, difference between means=2.161±1.069%, 95% CI=-4.616 to 0.2938] between YFP [lower 95% CI=0.1860, higher 95% CI=5.052] and LTD groups [lower 95% CI=-0.2284, higher 95% CI =1.144]). (N) Quantification of attack latency on the sixth day against a CD1 male (n=8 mice per group, two-tailed unpaired *t*-test, *P* = 0.0328 [observed power=0.985, Cohen’s D=1.0431, difference between means=227.00±82.23 sec, 95% CI =25.81 to 428.2] between YFP [lower 95% CI=67.01, higher 95% CI=477.9] and LTP groups [lower 95% CI=4.196, higher 95% CI=86.63], two-sided Mann–Whitney U test, *P* > 0.9999 between YFP and LTD groups). (O) Quantification of close investigation on the sixth day against a CD1 male (n=8 mice per group, two-tailed unpaired *t*-test, *P* = 0.6973 between YFP and LTP groups, *P* = 0.6158 between YFP and LTD groups). ns; not significant, \**P* < 0.05, \*\**P* < 0.01, \*\*\**P* < 0.001, \*\*\*\**P* < 0.0001. In box-and-whisker plots, center lines indicate medians, box edges represent the interquartile range, and whiskers extend to the minimal and maximal values.

The 5cdRI test was used to investigate the possible influence of LTP and LTD on aggression training. In one experiment, to determine whether LTP could facilitate aggression training, the LTP induction protocol was delivered at the end of each RI trial in both the control (YFP/mCherry-expressing) and LTP groups. In a separate experiment, to determine whether LTP was necessary for aggression training, the LTD induction protocol was delivered at the end of each RI trial in the control and LTD groups (Fig. 5B); application of LTD is expected to override any LTP that may have occurred (80). As in Fig. 1A, smaller size BALB/c intruders were introduced to Esr1-Cre residents, in which the LTP/LTD protocols were optogenetically delivered (Fig. 5C). Aggression levels were recorded and analyzed on each day of the 5cdRI (i.e., 24 hrs following the previous LTP or LTD manipulation, with the exception of day one).

Applying LTP or LTD induction protocols *in vivo* facilitated or diminished the behavioral effect of aggression training, respectively, as quantified by aggression/trial (% of the total trial duration occupied by aggressive behavior) and attack latency (Fig. 5D-K). Interestingly, although LTP was found to enhance the behavioral effect of aggression training on the second and third day of the 5cdRI assay, it did not lead to ever-increasing aggression levels; rather the effect plateaued on days four and five, at a level similar to the control group (see also Fig. 1C), suggesting an effect to accelerate learning. In contrast, LTD had a profound inhibitory effect on aggression training, leading to similar aggression levels between day one and day five of the 5cdRI test (Fig. 5D, two-tailed paired *t*-test, *P* = 0.0592 between day one and day five in the LTD group).

We investigated next whether the observation that control and LTP-induced groups expressed similar levels of aggressive behavior following training day three is due to a “ceiling effect” in the aggression training paradigm. To do this, we performed further tests following completion of the 5cdRI training routine. On day six, mouse social behavior was tested in a novel arena against a CD1 male conspecific of larger size (Fig. 5B, 5C), under which condition aggressive resident mice are less likely to attack (81). We reasoned that LTP mice that reached “ceiling” levels of aggression in the 5cdRI assays using conventional, smaller subordinate intruders might nevertheless show higher aggression under these sub-optimal conditions.

Indeed, under these conditions, the 5cdRI/LTP-treated mouse group exhibited higher aggression levels than any other tested group, while the control and LTD groups expressed similar aggression levels (Fig. 5L-O). This finding suggests that hypothalamic LTP expressed by VMHvl^Esr1^ neurons can facilitate aggression under modified conditions where resident aggressiveness is behaviorally reduced, relative to that typically detected in our conventional RI assay.

Together these experiments demonstrate a potential role for LTP and LTD in AGG mice. We next investigated the basis for individual differences in aggression training among genetically identical mice, by asking whether we could identify any experimental intervention that would allow aggression and/or hypothalamic LTP to be expressed in NON mice.

### Testosterone enables the expression of aggression and hypothalamic LTP in NON mice

Levels of testosterone (T) have been suggested to correlate with aggression and dominance in numerous species (82-90), while administration of T following castration or ovariectomy has been shown to restore aggression to the animal’s behavioral repertoire (91-96). As T levels are subject to epigenetic influences (97), we sought to determine whether individual differences in levels of the hormone were detectable among genetically identical, inbred C57BL/6N mice, and if so whether they correlated with and were responsible for individual differences in the capacity to undergo aggression training.

To investigate whether serum T levels differ between NON and AGG mice, we collected blood samples at different time points of the 5cdRI test (Fig. 6A-D). This experiment revealed that prior to the experience of aggression, a small but statistically significant (*P* < .05) difference in serum T is present between NON and AGG mice (Fig. 6B). This difference between the two groups was further accentuated following aggression training (Fig. 6C). This is because serum T levels remained unaltered following aggression training in NON mice, whereas they increased in AGG mice following training (Fig. 6D). Interestingly the increase in serum T in AGG mice occurred in the first three days, and was not further accentuated through additional aggression training (Fig. 6D). Together, these data reveal a correlation between individual differences in T and the ability to respond to aggression training in NON vs. AGG mice, as well as between levels of aggressiveness and T levels in AGG mice during training.

**Fig. 6.**
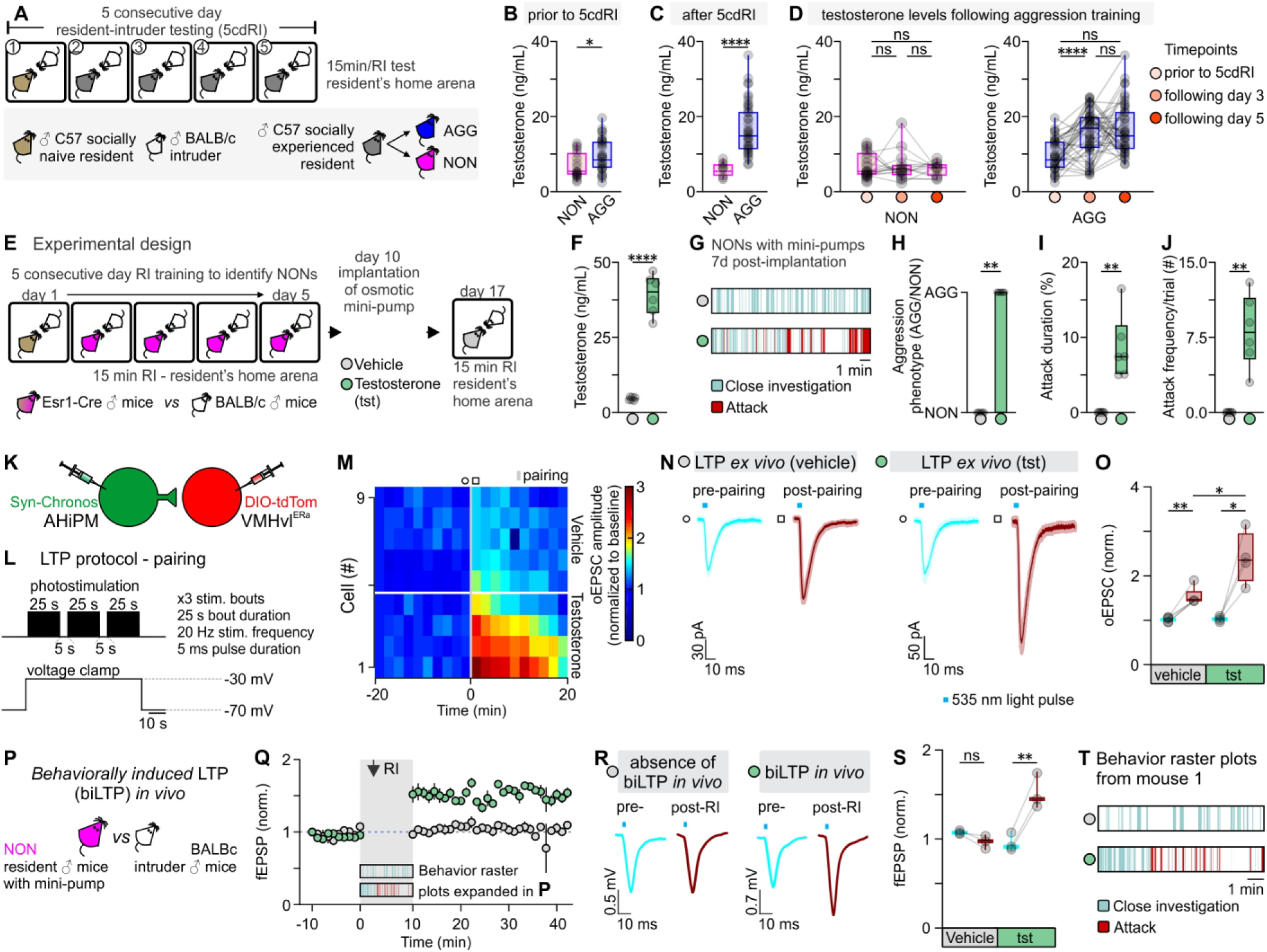
Testosterone administration leads to the expression of hypothalamic LTP and aggression in previously non-aggressive males. (A) Schematic of the experimental design used to identify aggressive (AGG) and non-aggressive (NON) males, from which tails blood samples were collected for quantification of serum testosterone levels. (B) Serum testosterone levels in NON *vs* AGG mice prior to any aggression experience (n=24-36 samples per group, two-sided Mann–Whitney U test, *P* = 0.0203 between NON and AGG groups). Mice were assigned as NON or AGG, according to whether they expressed aggression on the 1^st^ day of the 5cdRI test. (C) Serum testosterone levels in NON *vs* AGG mice after completion of the 5cdRI test (n=14-46 samples per group, two-tailed unpaired *t*-test, *P* < 0.0001 between NON and AGG groups). Mice that did not express any aggression/attack behavior throughout the 5cdRI test they were assigned to the NON group. All other mice they were included in the AGG group. (D) Left – Quantification of serum testosterone levels in NON mice throughout the 5cdRI test (n=14-24 samples per group, Kruskal-Wallis one-way ANOVA with Dunn’s post hoc test, *P* > 0.9999 between *prior to 5cdRI* and *following day 3*, *P* > 0.9999 between *prior to 5cdRI* and *following day 5*, and *P* > 0.9999 between *following day 3* and *following day 5* groups). Right – Quantification of serum testosterone levels in AGG mice throughout the 5cdRI test (n=36-46 samples per group, one-way ANOVA with Tukey’s test, *P* < 0.0001 between *prior to 5cdRI* and *following day 3*, *P* < 0.0001 between *prior to 5cdRI* and *following day 5*, and *P* = 0.7060 between *following day 3* and *following day 5* groups). (E) Schematic of the experimental design used to identify NON mice, and perform subcutaneous testosterone mini-pump implantation. (F) Serum testosterone levels in control vs. testosterone-treated mice (n=6 mice per group, two-tailed unpaired *t*-test, *P* < 0.0001 between vehicle and testosterone). (G) Representative behavior raster plots of vehicle *vs* testosterone-treated mice. (H) Quantification of the number of mice that switched aggression phenotype following vehicle *vs* testosterone administration (n=0/6 in the vehicle-treated group *vs* n=6/6 in the testosterone treated group, two-sided Mann–Whitney U test, *P* = 0.0022 between vehicle and testosterone). (I) Quantification of attack duration (n=6 mice per group, two-sided Mann–Whitney U test, *P* = 0.0022 between vehicle and testosterone). (J) Quantification of attack frequency (# attacks/trial; n=6 mice per group, two-sided Mann–Whitney U test, *P* = 0.0022 between vehicle and testosterone). (K) Schematic of the experimental design used to study the induction and modulation of LTP by testosterone in the AHiPM➔VMHvl synapse in brain slices from NON mice. Note that slices were taken from animals that received T injections, but no behavioral training or other social experience. (L) Schematic of the LTP induction protocol, utilizing simultaneous photostimulation of the AHiPM terminals in VMHvl through the opsin Chronos and depolarization of the VMHvl^Esr1^ neuron through voltage clamp at −30 mV. (M) Heat map illustrating the magnitude of LTP induction in VMHvl^Esr1^ neurons from vehicle- *vs* testosterone-treated mice. (N) Average current immediately prior to and following the induction of LTP in vehicle *vs* testosterone conditions (light color envelope is the standard error). (O) Quantification of the optically evoked excitatory post-synaptic current (oEPSC), prior to and following the induction of LTP in vehicle *vs* testosterone conditions (pre-[lower 95% CI= 0.9508, higher 95% CI=1.084] *vs* post-[lower 95% CI=1.274, higher 95% CI=1.786] pairing in vehicle conditions, n=5 cells from 3 mice, two-tailed paired *t*-test, *P* = 0.0038 [observed power=0.992, Cohen’s D=2.704, difference between means=0.5124±0.0847, 95% CI=0.2771 to 0.7478], pre-[lower 95% CI=0.9349, higher 95% CI=1.120] *vs* post-[lower 95% CI=1.456, higher 95% CI=3.334] pairing in testosterone conditions, n=4 cells from 3 mice, two-tailed paired *t*-test, *P* = 0.0209 [observed power=0.889, Cohen’s D=2.232, difference between means=1.368±0.3063, 95% CI=0.3926 to 2.342], post-pairing in vehicle [lower 95% CI=1.274, higher 95% CI=1.786] *vs* testosterone [lower 95% CI=1.456, higher 95% CI=3.334] conditions, n=4-5 cells from 6 mice, two-tailed unpaired *t*-test, *P* = 0.0174 [observed power=0.932, Cohen’s D=0.6388, difference between means=0.8652±0.2974, 95% CI=0.2044 to 1.526]). (P) Schematic of the experimental design used to trigger and record behaviorally induced LTP *in vivo* in NONs. (Q) Field EPSP amplitude (fEPSP) over time, prior to and following social behavior in the resident intruder assay, in vehicle- *vs* testosterone-treated NON mice (average fEPSP from n=3 mice per group). (R) Average fEPSP amplitude immediately prior to and following the expression of social behavior in the resident intruder assay, in vehicle- *vs* testosterone-treated NON mice. (S) Quantification of fEPSP amplitude, prior to and following the induction of LTP in vehicle *vs* testosterone conditions (pre-[lower 95% CI=1.027, higher 95% CI=1.123] *vs* post-[lower 95% CI=0.7907, higher 95% CI=1.143] pairing in vehicle conditions, n=3 mice, two-tailed paired *t*-test, *P* = 0.1020 [observed power=0.999, Cohen’s D=1.6667, difference between means=0.1081±0.0374, 95% CI=− 0.2692 to 0.05303], pre-[lower 95% CI=0.7055, higher 95% CI=1.209] *vs* post-[lower 95% CI=1.027, higher 95% CI=2.012] pairing in testosterone conditions, n=3 mice, two-tailed paired *t*-test, *P* = 0.0098 [observed power=0.786, Cohen’s D=5.7787, difference between means=0.5625±0.0562, 95% CI=0.3207 to 0.8043]). (T) Representative behavior raster plot of the same mouse treated with vehicle and 8 days after with testosterone and used for *in vivo* electrophysiology experiments. ns; not significant, \**P* < 0.05, \*\**P* < 0.01, \*\*\*\**P* < 0.0001. In box-and-whisker plots, center lines indicate medians, box edges represent the interquartile range, and whiskers extend to the minimal and maximal values. In bar graphs, data are expressed as mean ± s.e.m.

To test whether T levels are causally responsible for the difference in aggressiveness between NON and AGG mice, subcutaneous osmotic mini-pumps containing T or vehicle were implanted in NON mice (Fig. 6E). The serum T levels in NONs measured at seven days post T mini-pump implantation were significantly higher than (but within the upper quartile of) the endogenous T levels measured in AGG mice following completion of the 5cdRI training (Fig. 6D, F, serum T in NONs following T administration through mini-pump 39.22±2.73 ng/mL, serum T in AGGs following aggression training 16.61±1.01 ng/mL, n=6 and 46 mice respectively, two-sided Mann–Whitney U test, *P* < 0.0001). Strikingly, the administration of exogenous T induced aggression in all NON mice tested (Fig. 6G-J). We next investigated whether the emergence of aggression through T administration correlated with the expression of LTP. In acute VMHvl slices from vehicle *vs* T-treated NON mice, we investigated the induction and expression of LTP using *ex vivo* recordings. Using the same Hebbian induction protocol, stronger LTP could be elicited from VMHvl^Esr1^ neurons recorded in slices from T-treated NON mice, in comparison to those from vehicle-treated NON mice (Fig. 6K-O). Thus, T implants facilitate LTP induction *ex vivo* in NON mice.

An important remaining question, however, was whether LTP was expressed in VMHvl^Esr1^ neurons *in vivo*, following aggression training in T-treated NON mice. To address this question, we used the design previously described in Fig. 4H, N, in which AHiPM was transduced with Chronos, while VMHvl^Esr1^ cells were transduced with ChrimsonR. A novel BALB/c, small size male intruder was introduced into the NON’s home cage (Fig. 6P). In vehicle-treated mice, social interactions with intruder mice, but no aggression, were observed, and LTP did not occur *in vivo,* as measured by fEPSP recordings in response to optogenetic stimulation of Chronos-expressing AHiPM terminals (Fig. 6Q-T). However, T administration through subcutaneous osmotic mini-pumps led to the expression of both aggressive behavior and *in vivo* behaviorally induced LTP, in NON mice (Fig. 6Q-T).

These findings suggest that individual differences in serum T are responsible, at least in part, for individual differences in the capacity for aggression training amongst inbred mice. Elevation of serum T in NON mice can restore susceptibility to aggression training, as well as the capacity to express strong LTP at AHiPM➔VMHvl^Esr1^ synapses (both *ex vivo* and *in vivo* following aggression training). This observation further strengthens the correlation between the ability to respond positively to aggression training, and the expression of LTP. However, it does not distinguish whether the enhanced LTP in NON mice is directly caused by T treatment, or rather is an indirect effect of the increased aggression promoted by the hormone implants.

## Discussion

A prevailing view in neuroscience is that anatomically distinct neural systems mediate innate *vs*. learned behaviors: the former are thought to be processed by “labelled lines,” developmentally hard-wired circuits in evolutionarily ancient, deep subcortical structures such as the hypothalamus (13, 98, 99); in contrast, the latter engage synaptic plasticity mechanisms in more recently evolved brain structures, such as the neocortex and hippocampus. Reflecting this view, in the mammalian brain the vast majority of studies of synaptic plasticity mechanisms, such as LTP and LTD, have been performed in the latter structures (as well as in the cerebellum (100-102). Whether such mechanisms also operate in deep subcortical structures, and if so, what types of behavioral plasticity (if any) they might subserve, has remained unclear. However, this knowledge gap reflects an absence of evidence, more than evidence of absence.

Here we have identified and deconstructed the neural substrate and physiological mechanism underlying a form of experience-dependent plasticity in aggression, a prototypic innate social behavior. We show that a training paradigm that increases aggressiveness via repeated successful agonistic encounters is correlated with, enhanced by and dependent upon, LTP operating at a glutamatergic synapse on a population of hypothalamic Esr1^+^ neurons that mediates innate aggressive behavior (47). The plasticity observed at AHiPM➔VMHvl^Esr1^ synapses likely has both post- and pre-synaptic components, as suggested by an increase in the AMPAR/NMDAR ratio (54-56) following aggression training (Fig. 2J, K), and by the differential responses of VMHvl^Esr1^ neurons to trains of pre-synaptic stimuli (103, 104) (Fig. 2L, M), respectively. Surprisingly, the form of hypothalamic LTP studied here does not exhibit “occlusion,” phenomenon observed in studies of hippocampal or amygdalar LTP (105, 106), in which following *in vivo* behavioral induction of LTP in the synaptic population of interest, the magnitude of LTP that can be induced subsequently *ex vivo* is markedly decreased (Fig. 4D, K). Similarly, we do not observe the related phenomenon in which prior *in vivo* LTP can enhance the extent of LTD that can be induced *ex vivo* in slices from such animals. The reason(s) for the failure to observe these phenomena are not clear, and will require further studies to elucidate. There are a number of effects, however, which could account for these observations. Firstly, it is possible that the proportion of synapses modified by the *in vivo* social experience was small compared to the synapses being sampled in the slice. Another possibility is that the synapses being assayed in the slice are a different population than the ones modified *in vivo*, or lastly that new synapses were formed by the *in vivo* experience and they are the ones primarily contributing to the LTP and LTD being measured *in vitro*. This last possibility is of particular interest, given that - as presented in Fig. 3, an increase in spine density occurs in VMHvl^Esr1^ neurons of AGG mice.

The data on LTP presented here, blur the distinction between neural circuits mediating learned vs. innate behaviors, and reinforce the concept of “learned innate behavior,” in which synaptic plasticity within developmentally hardwired circuits can function to modify the strength of an instinctive behavior in response to social experience. An example of the latter in an invertebrate is the post-mating response in *Drosophila,* a form of memory in which female sexual receptivity is inhibited following mating (107-109). Interestingly, recent studies have blurred the classic distinction between the innate and adaptive immune systems as well (110, 111).

This idea notwithstanding, more complex forms of learning, such as classical or operant conditioning, may utilize circuits that are parallel to those that mediate innate forms of the modified behavior, as shown in the case of conditioned *vs*. unconditioned fear (112-114). In this context, it is worth noting that mice can learn an instrumental, operant response using successful aggressive encounters as a reinforcer (115), and that performance of this instrumental task is facilitated by optogenetic activation of VMHvl neurons (116). The neural substrates and synaptic mechanisms underlying this operant conditioning remain to be elucidated, although the nucleus accumbens-based reward system has been implicated in recent studies (117).

Aggressiveness can be enhanced not only by repeated successful agonistic encounters, as shown here, but also by prior mating experience (118). Recently, we showed that as little as 30 minutes of free social interaction with a female was sufficient to transform a socially naive mouse into an AGG mouse within 24 hrs of the interaction (119). This effect was associated with a change in the neural representation of male *vs*. female conspecifics among VMHvl^Esr1^ neurons, from partially overlapping to largely non-overlapping (119). Whether this change in neural population coding involves synaptic plasticity within VMHvl, or is inherited from upstream structures, such as the MeA (120), remains to be determined. In other studies, we have shown that the effect of social isolation stress to promote aggression in non-sexually experienced males is mediated by the neuropeptide Neurokinin B (NkB) and its receptor Nk3R, acting in the dorso-medial hypothalamus (DMH) (121). The relationship of this form of experience-dependent plasticity to VMHvl^Esr1^ neuronal activity is currently unknown.

Our current findings also provide insights into individual differences in the ability of genetically identical animals to respond to “aggression training”. Firstly, we show here that several physiological parameters in AGG mice are different from those in socially naïve mice. These include elevated baseline VMHvl^Esr1^ neuron activity (Fig. 1G-J), increased spontaneous excitatory input onto VMHvl^Esr1^ neurons (Fig. 2A-C), increased AMPA/NMDA ratio at AHiPM➔VMHvl^Esr1^ synapses (Fig. 2I-K) and altered synaptic integration properties (Fig. L, M). By contrast, in NON mice the spontaneous inhibitory inputs to VMHvl^Esr1^ neurons are increased, relative to socially naïve mice (Fig. S1). In addition, NON mice exhibit shorter lasting LTP and longer lasting LTD than are observed in AGG mice (Fig. 4D, G). Whether increased LTD is sufficient to account for the failure of NON mice to respond to aggression training is not yet clear. Another possibility, suggested by the increased spontaneous IPSCs, is that VMHvl receives stronger inhibitory input from GABAergic neurons in NON mice. While there are very few GABAergic neurons within VMHvl itself (122), VMHvl receives strong inhibitory input from the neighboring tuberal (TU) region. It is possible that the lack of aggression in NON mice reflects potentiation of these TU GABAergic neurons. Whether these TU neurons receive feed-forward input from VMHvl^Esr1^ neurons, or from another source, is not known. The synaptic mechanisms responsible for the lack of aggression in NON mice will clearly require further investigation.

We also find that NON mice – unsusceptible to aggression training, have low levels of circulating T in comparison to AGG mice, and experimental administration of supplemental T can restore the capacity for “aggression learning” in such animals. While the permissive role of T in promoting male aggressiveness is well-established (82, 86, 123-126), our studies provide new insight into the neurophysiological mechanisms that may mediate this effect in the context of aggression training. Specifically, we observe that NON animals can only express LTP *in vivo* following administration of exogenous T. Although LTP can be induced optogenetically *ex vivo* in slices from control NON animals, LTP in slices from T-implanted NON animals exhibited higher-amplitude and persistence. Moreover, in AGG mice levels of T increased during aggression training. This correlation suggests either that T acts directly to enhance LTP at this synapse, which in turn promotes aggression, or that T acts indirectly, by promoting aggressive behavior which in turn enhances LTP (Fig. S5). Whether T directly influences synaptic plasticity, and if so the underlying molecular mechanisms involved, as well as the mechanistic basis of individual differences in T levels, are interesting topics for future study.

Our experiments have focused on a specific glutamatergic input to VMHvl^Esr1^ neurons which have a causal role in aggression. In addition to our finding, recent work reported that VMHvl-projecting Vglut^+^ neurons in the AHiPM exhibited elevated c-fos expression following both social investigation and attack, while chemogenectic silencing of AHiPM neurons inhibited attack (51). VMHvl^Esr1^ neurons receive inputs from neurons in over 30 different structures (50), raising the question of whether other inputs to these cells also display plasticity. Indeed, recently published work has identified synaptic plasticity promoted by foot-shock stress in a medial amygdala projection that primarily targets the central part of VMH (VMHc) (127). Although a causal role in promoting aggression was not directly demonstrated for this input, and the mechanism of potentiation was not established, plasticity at this synapse may regulate stress-induced aggression (127). The present study demonstrates that AHiPM➔ VMHvl^Esr1^ synapses can undergo Hebbian LTP, and that potentiation of these synapses occurs during social experience that enhances offensive aggression (47, 119). Together, these data suggest that VMHvl likely provides a substrate in which aggression plasticity can occur at multiple synaptic inputs, each of which may play distinct roles in physiology and/or behavior. Our results also reveal striking effects of aggression training on dendritic spine morphology among VMHvl^Esr1^ neurons, although we cannot be certain whether the secondary dendritic branches where we observe this phenomenon receive synaptic input from AHiPM. Other recent studies have identified structural plasticity among VMHvl^PR^-derived axons innervating hypothalamic targets in females, which are mediated by changes in sex steroids during estrus (128). The present work, together with these other studies, begins to provide a view of the acute and dynamic changes that can occur through experience and/or hormonal modulation, in a brain node that controls innate social behaviors.

Historically, synaptic plasticity mechanisms – and in particular LTP, have been investigated predominantly in hippocampal circuits that mediate spatial learning (129-132), or in thalamo-amygdalar circuits that mediate Pavlovian associative conditioning (133-136). Both systems emphasize the role of LTP in allowing flexible neural circuits to mediate adaptive responses on fast time-scales, as expected for the recently evolved brain regions in which they operate. By contrast, studies of the hypothalamus have focused primarily on identifying circuits that mediate evolutionarily ancient, innate survival behaviors, with the expectation that such circuits would be comprised predominantly of relatively stable, hard-wired synaptic connections (13, 98, 99). Our results and other data suggest a reconsideration of this prevailing view of hypothalamic pathways as ‘hard-wired’ neural circuits. They suggest, moreover, that further investigation of synaptic plasticity mechanisms within neural pathways that control evolutionarily selected, robust survival behaviors, may yield new insights into both the physiological and hormonal regulation of such mechanisms, as well as the forms of behavioral plasticity that they ultimately subserve.

## Materials and Methods

All experimental procedures involving the use of live animals or their tissues were carried out in accordance with the NIH guidelines and approved by the Institutional Animal Care and Use Committee and the Institutional Biosafety Committee at the California Institute of Technology (Caltech). *Esr1^Cre/^*^+^ knock-in mice (47) backcrossed into the C57BL/6N background (>N10) were bred at Caltech. The *Esr1^Cre/^*^+^ knock-in mouse line is available from the Jackson Laboratory (Stock no. 017911). Heterozygous *Esr1^Cre/^*^+^ mice were used for all experiments and were genotyped by PCR analysis of tail DNA. Mice used as residents (see five consecutive-day resident–intruder assay) were individually housed. All wild-type mice used as intruders in resident–intruder assays and for behavioral experiments were of the BALB/cAnNCrl or Crl:CD1 (ICR) strain (Charles River Laboratories). Health status was normal for all animals. Antibodies, compounds, and the experimental procedures with the coordinates of all injection sites are described in *SI Appendix*.

## Data Availability

All data discussed in the paper are available in the main text and *SI Appendix*. We used standard MATLAB functions and publicly available software indicated in the manuscript for analysis.

## Acknowledgments

The authors thank Dr. B. Weissbourd and Dr. L. Li for advice on experiments, Dr. Y. Ouadah and Dr. P. Williams for advice during writing of the manuscript, X. Da and J.S. Chang for technical assistance, X. Da and C. Chiu for laboratory management and G. Mancuso for administrative support. Members of the Anderson laboratory are thanked for discussion during the preparation of this manuscript. Confocal imaging was performed in the Biological Imaging Facility, with the support of the Caltech Beckman Institute and the Arnold and Mabel Beckman Foundation. This study was supported by NIH Grant R01 MH070053 to D.J.A., and the EMBO ALTF 736-2018 postdoctoral fellowship to S.S. D.J.A. is an investigator of the Howard Hughes Medical Institute.

## Supplementary Information Text

### Extended materials and methods

#### Animals

All mice were housed in ventilated micro-isolator cages in a temperature-controlled environment (median temperature 23 °C), under a reversed 12h dark-light cycle, and had *ad libitum* access to food and water. Mouse cages were changed weekly on a fixed day on which experiments were not performed.

#### Brain slice electrophysiology

Acute mouse brain slices were prepared. Slices were cut on a vibratome (Leica VT1000S) to 300 μm thickness and continuously perfused with oxygenated aCSF containing (in millimolar): NaCl (127), KCl (2.0), NaH_2_PO_4_ (1.2), NaHCO_3_ (26), MgCl_2_ (1.3), CaCl_2_ (2.4), and D-glucose (10). See also Table S1. Whole-cell current- and voltage-clamp recordings were performed with micropipettes filled with intracellular solution containing (in millimolar), K-gluconate (140), KCl (10), HEPES (10), EGTA (10), and Na_2_ATP (2) or Cesium methanesulfonate (140), KCl (10), HEPES (10), EGTA (10), and Na_2_ATP (2) (pH 7.3 with KOH). Recordings were performed using a Multiclamp 700B amplifier, a DigiData 1440 digitizer, and pClamp 11 software (Molecular Devices). Slow and fast capacitative components were semi-automatically compensated. Access resistance was monitored throughout the experiments, and neurons in which the series resistance exceeded 15 MΩ or changed ≥20% were excluded from the statistics. The liquid junction potential was 9.7 mV and not compensated. The recorded current was sampled at 20 kHz. Baseline recordings of EPSCs, IPSCs and optogenetically-evoked synaptic currents were performed in normal aCSF conditions and in the absence of GABA and NMDA receptor blockers. Spontaneous excitatory currents were sampled at the reversal of Cl^-^ (V_HOLD_=-70 mV), and spontaneous inhibitory currents were sampled at the reversal of fast excitatory neurotransmission (V_HOLD_=0 mV). All recordings were performed at near-physiological temperature (33±1°C). Reagents used in slice electrophysiology experiments; Neurobiotin^TM^ tracer (Vector laboratories) was used in combination with Streptavidin conjugated to Alexa Fluor 647. MATLAB and OriginPro9 were used for electrophysiological data analysis. CNQX (10 μM), D-AP5 (25 μM), TTX (500 nM), and 4-AP (100 mM) were bath applied to block excitatory transmission and to test if optogenetically evoked responses are monosynaptic (137). All drugs were pre-applied for 5 min in the slice chamber prior to data acquisition.

#### Brain slice Ca^2+^ imaging

The spontaneous activity of mouse VMHvl^Esr1^ neurons was monitored by imaging fluorescence changes of the jGCaMP7s biosensor, using a CCD camera (Evolve^®^ 512, Photometrics), mounted on an Olympus BX51WI microscope. Recordings were 5 min in duration. As a subpopulation of VMHvl^Esr1^ neurons expresses T-type Ca^2+^ channels (*unpublished data)*, the Ca^2+^ transients reported in Fig. 1 likely reflect both action potentials and subthreshold synaptic potentials. A 60x water-dipping objective was used to focus on VMHvl. Ca^2+^ imaging analysis was performed using the MIN1PIPE one-photon based calcium imaging signal extraction pipeline (138), in combination with custom-written MATLAB routines.

#### Cell filling and reconstruction

Mouse *Esr1*^+^ VMHvl neurons were recorded in whole-cell mode with intracellular pipette solution as above, with the addition of 0.2% neurobiotin. After recording, slices were placed in fixative (4% paraformaldehyde/0.16% picric acid), washed in PBS and incubated at 4°C for 72h in a solution containing streptavidin conjugated to Alexa Fluor 647. After extensive washing, slices were mounted with 2.5% DABCO in glycerol. VMHvl^Esr1^ neuron identity of all filled cells was confirmed with colocalization studies of viral-induced tdTomato expression.

#### *Ex vivo* optogenetics

Photostimulation during slice whole-cell recordings was performed via a 3.4 watt 535 nm LED mounted on the microscope fluorescence light source and delivered through the 60X objective’s lens. Photostimulation was controlled via the analog outputs of a DigiData 1440A, enabling control over the duration and intensity. The photostimulation diameter through the objective lens was ∼310 μm with illumination intensity typically scaled to 0.35 mW/mm^2^.

#### *In vivo* optogenetics

Subjects were coupled via a ferrule patch cord to a ferrule on the head of the mouse using a zirconia split sleeve (Doric Lenses). Ferrules and fiber-optic patch cords were purchased from Thorlabs and Doric Lenses, respectively. The optical fiber was connected to THORLABS fiber-coupled LED (M530F2, 9.6 mW) via a fiber-optic rotary joint (FRJ_1x1_FC-FC, Doric Lenses) to avoid twisting of the cable caused by the animal’s movement. Prior to a testing session, following the coupling of the patch cords with the optic fiber ferrules, *Esr1^Cre/^*^+^ animals were given 10 min to acclimate in their home cage in the absence of an intruder. The frequency and duration of photostimulation were controlled using the programmable train generator Pulse Pal (139). Light power was controlled by dialing an analog knob on the LED driver (T-Cube^TM^ LED Driver with Trigger Mode, Thorlabs, LEDD1B). Light power was measured from the tip of the ferrule in the patch cord at different laser output settings, using an optical power energy meter and a photodiode power sensor (Thorlabs, PM100D, and S130VC). Light power was dialed at 0.5 mW at the fiber tip. Upon identification of the fiber placement coordinates in brain tissue slides, irradiance (light intensity) was calculated using the brain tissue light transmission calculator based on (http://www.stanford.edu/group/dlab/cgi-bin/graph/chart.php) using laser power measured at the tip and the distance from the tip to the target brain region measured by histology. Animals showing no detectable viral expression in the target region and/or ectopic fiber placement were excluded from the analysis.

#### *In vivo* electrophysiology

*In vivo* electrophysiology recordings were performed in freely moving mice, using chronic silicon probe implants. All extracellular recordings were conducted in the left VMHvl, and all mice included in the present study were validated using the following criteria: identification of the lipophilic dye (DiD) tract targeting VMHvl, phototagging of VMHvl^Esr1^ neurons, and photostimulation-evoked low-latency attack against a conspecific through optrode mediated VMHvl^Esr1^ neuron photoactivation. Recordings were performed using an optrode based on the A1x32-Poly2-10mm-50s-177 NeuroNexus probe and a 100 μm optic fiber placed along the probe’s shank terminating 50 μm above the probe’s first recording sites. Photostimulation was delivered using fiber-coupled Thorlabs LEDs (M530F2, 9.6 mW for LTP/LTD studies, and M617F2, 13.2 mW for phototagging), and light power was dialed at 0.33 mW at the optrode’s fiber tip. The probe was implanted 200 μm above the intended recording site, and using the NeuroNexus OH32LP oDrive was lowered over a period of four days to the target coordinates (lowering by 50 μm/day). Only channels that showed photo-responses in the local field potential were used for LFP analysis. Recordings were performed using the Open Ephys acquisition board with a sampling rate of 30 kHz, the Open Ephys I/O board, and the Open Ephys GUI (140). The LTP signal was obtained by applying low pass-filtering with a cut-off at 100 Hz on the raw voltage traces.

#### Immunohistochemistry

Mice were anesthetized with ketamine (KetaVed, VEDCO) and xylazine (AnaSed, NDC 59399-110-20), then transcardially perfused with 20 mL of ice-cold fixative. Whole brains were dissected, immersed in ice-cold fixative for 90 min then stored in 0.1M PBS (pH 7.4) containing 20% sucrose, 0.02% bacitracin and 0.01% sodium azide for three days, before freezing with dry ice. Coronal sections were cut at a thickness of 14 μm on a cryostat (Microm, Walldorf) and thaw-mounted onto gelatine-coated glass slides. For GFP staining, brain sections were incubated overnight at 4°C using a chicken anti-GFP antibody (Aves Labs, Inc., GFP-1010) at 1:500 dilution. For tdTomato staining brain sections were incubated overnight at 4°C using a rabbit anti-DsRed antibody (Takara, 632392) at 1:500 dilution. Primary antibody incubation was followed by Alexa-488-conjugated goat anti-chicken secondary antisera (1:500; Invitrogen), and/or Alexa-568-conjugated donkey anti-rabbit secondary antisera (1:500; Invitrogen). DAPI solution (1mg/mL) was used at 1:10000 dilution. For further details on reagents, see also Table S1.

#### Confocal microscopy

Brain slices were imaged by confocal microscopy (Zeiss, LSM 800). Brain areas were determined according to their anatomy using Paxinos and Franklin Brain Atlas (141).

For cell reconstructions, each entire neurobiotin-filled neuron was acquired at 63X (NA = 1.4), 1 µm step size using a Zeiss LSM880 confocal microscope. Imaris 9.3 (Bitplane) was used to visualize the topology of the dendritic tree and the centrifugal branch ordering method was chosen to sort dendrites, assigning order 1 to the root. 2^nd^ order dendrites were then selected for further imaging acquisition to perform spine quantification. 70-90 µm-long dendritic segments were acquired at 63X (NA = 1.43), 0.1 µm step size and 0.06×0.06 pixel-size using Airy-scan detector at the LSM880. Two segments were acquired for dendrites longer that 200 µm.

For spine quantification, images of dendritic segments were rendered in Imaris using the *Blend* algorithm and the *Filament* module was used to reconstruct dendrites and spines. Specifically, the auto-path method was chosen and thinnest spine diameter (between 1.5 and 2 µm), maximal distance from the dendrite (between 3 and 8 µm) and fluorescence intensity threshold were defined in every single dendrite to detect spines. The statistics module in Imaris was used to extract spine density values. Three to six segments per neuron were quantified and values were averaged.

#### Tail-tip whole blood sampling

Whole blood samples of 40-70 μL were collected from the lateral tail vein of restrained mice (142). Only blood samples acquired within 2 min post-restraining were used for hormone measurements, and the subjects were then returned to their home cage. Briefly, the rodent’s tail was immersed for 30 sec in 40°C water to dilate the tail blood vessels. Immediately after, a 23G needle was used to puncture the lateral tail vein, and whole blood was collected. Bleeding was stopped via applying gentle pressure to the tail at the level of the puncture with surgical cleaning tissue, and 2% chlorhexidine antiseptic solution was used for tail disinfection at the end of the procedure. Blood samples were refrigerated at 4°C for 30 min and then centrifuged at 4^ο^C at 2000 RCF. Following centrifugation, serum was collected and was frozen at −80°C for a maximal period of 2 months prior to performing ELISA measurements. All blood samples were acquired during the dark phase of the 12h/12h light/dark cycle. For further details on reagents, see also Table S1.

#### Testosterone ELISA

96-well plates were used in a ready-to-use kit for testosterone ELISA (R&D systems – Catalog number KGE010). Linear regression was used to fit the optical densities for the standard curve *vs* the concentration. The standard curve range for corticosterone was 300 to 100000 pg/mL. Concentrations were calculated from the optical density at 450 nm of each sample. Appropriate sample dilutions were carried out to maintain detection in the linear part of the standard curve and typically involved 1 to 10 for mouse serum samples. Data acquired from the performed ELISAs are presented as absolute values. Differences between groups were identified by Student’s *t-*test or ANOVA.

#### Viral vectors

For *ex vivo* Ca^2+^ imaging studies of VMHvl neurons, *Esr1Cre/*+ male mice were injected in VMHvl with 200 nL of AAV9-Syn-FLEX-jGCaMP7s-WPRE (addgene 104491-AAV9) 5.3 × 10^12^ genomic copies per mL. For *ex vivo* optogenetic studies, *Esr1Cre/*+ male mice were injected in VMHvl with 200 nL of AAV9-FLEX-tdTomato (addgene 28306-AAV9) 4.2 × 10^12^ genomic copies per mL and in AHiPM with 100 nL of AAV5-Syn-Chronos-GFP (addgene 59170-AAV5) 3.7 × 10^12^ genomic copies per mL. For *in vivo* optogenetic and electrophysiology experiments, *Esr1Cre/*+ male mice were injected in VMHvl with 100 nL of AAV5-Syn-FLEX-rc[ChrimsonR-tdTomato] (addgene 62723-AAV5) 4.1 × 10^12^ genomic copies per mL and in AHiPM with 100 nL of AAV5-Syn-Chronos-GFP (addgene 59170-AAV5) 3.7 × 10^12^ genomic copies per mL. Control groups were injected in VMHvl with 100 nL of AAV9-FLEX-tdTomato (addgene 28306-AAV9) 4.2 × 10^12^ genomic copies per mL and in AHiPM with 100 nL of AAV5-CAG-GFP (37825-AAV5) 5.9 × 10^12^ genomic copies per mL. For further details on reagents, see also Table S1.

#### Stereotactic surgery and viral gene transfer

Adult heterozygous *Esr1Cre/*+ males were single-housed for at least five days before undergoing surgical procedures and were operated on at 16–20 weeks of age. Mice were anesthetized using isoflurane (5% induction, 1–2% maintenance, in 95% oxygen) and placed in a stereotaxic frame (David Kopf Instruments). Body temperature was maintained using a heating pad. An incision was made to expose the skull for stereotaxic alignment using the inferior cerebral vein and the Bregma as vertical references. We based the coordinates for the craniotomy and stereotaxic injection of VMHvl on an anatomical magnetic resonance atlas of the mouse brain (AP: −4.68 mm; ML: ±0.78 mm; DV: −5.80 mm), as previously described (47). Virus suspension was injected using a pulled-glass capillary at a slow rate of 8–10 nL/min, 100 nl per injection site (Nanojector II, Drummond Scientific; Micro4 controller, World Precision Instruments). The glass capillary was withdrawn 10 min after the cessation of injection.

#### Osmotic mini-pumps

Testosterone was dissolved at 30 mg/ml in sesame oil and was administered for 2 weeks at a rate of 0.75 mg/hour via subcutaneous osmotic mini-pumps (Alzet, model 1002) (143-145). For further details on reagents, see also Table S1.

#### Social behavior assays

The aggression phenotype of animals defined as aggressive (AGG), or non-aggressive (NON) in the present study was based on the expression of aggressive behavior in the five consecutive day resident-intruder test (5cdRI). Animals that did not express any aggressive behavior in the 5cdRI were identified as NONs, while all AGGs expressed aggression in a minimum of three out of the five trials, with the majority expressing attack behavior in all five days. As described in Fig. 1, the 5cdRI composed of a 15 min social interaction test per day in the resident’s home arena, with socially naïve 4-5 month-old residents. Intruders were BALB/c males 2-3 months old and of lower weight/size. Three follow up tests were performed in the 5cdRI experimental design presented in Fig. 1, specifically, 2 weeks, 4 weeks and 12 weeks following the completion date of the 5cdRI assay. Note that only 15 out of a total of 106 aggressive mice, were used to quantify the effect of aggression training. This was based on the finding that following behavioral analysis of the first 15 mice used in the study, the power of the ANOVA test reached *P* < 0.0001. This suggested that including additional observations would not aid the power of the statistical test. In Fig. 5, following the 5cdRI, on day six a social interaction test was performed in a novel home-cage-sized arena. In addition to the C57 male, a male with a larger bodyweight/size CD-1 conspecific was introduced. The duration of this experiment was 15 min, following which both animals were returned to their home cage.

#### Statistics

No statistical methods were used to predetermine sample sizes but our sample sizes are similar to those reported in previous publications (35, 44, 47). Data met the assumptions of the statistical tests used and were tested for normality and variance. Normality was determined by D’Agostino–Pearson normality test. All *t*-tests and one-way ANOVAs were performed using GraphPad Prism software (Graphpad Software Inc.). Statistical significance was set at *P* < 0.05.

---

**Fig. S1.**
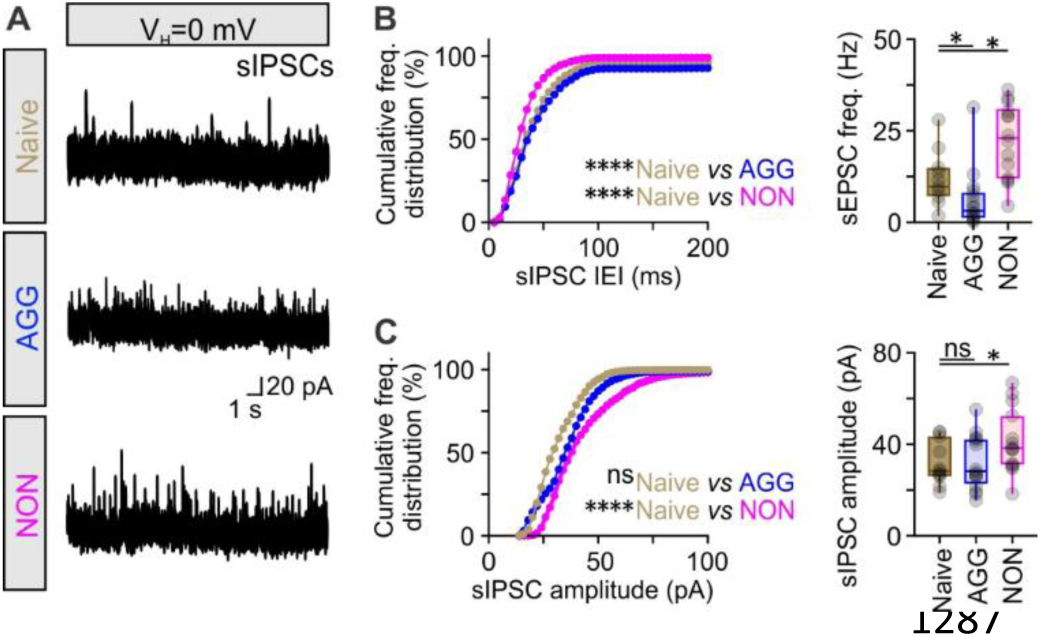
Presynaptic plasticity of inhibitory input in VMHvl^Esr1^ neurons of non-aggressive male mice. (A) Representative recordings of spontaneous inhibitory post-synaptic currents (sIPSCs) from VMHvl^Esr1^ neurons, from socially naive, aggressive (AGG) and non-aggressive (NON) mice. (B) Left – cumulative frequency distribution plot of sIPSC inter-event interval (IEI) in voltage-clamp recordings collected from VMHvl^Esr1^ neurons from socially naive, AGG and NON mice (n=11-14 VMHvl^Esr1^ neuron recording per group, collected from 8-10 mice per group, Kolmogorov-Smirnov test, *P* < 0.0001 between socially naive and AGG mice, *P* < 0.0001 between socially naive and NON mice). Right – comparison of sIPSC frequency in voltage-clamp recordings collected from VMHvl^Esr1^ neurons from socially naive, AGG and NON mice (n=11-14 VMHvl^Esr1^ neuron recording per group, collected from 8-10 mice per group, Kruskal-Wallis one-way ANOVA with uncorrected Dunn’s post hoc test, *P* = 0.0425 between socially naive and AGG mice, *P* = 0.0480 between socially naive and NON mice). (C) Left – cumulative frequency distribution plot of sIPSC amplitude in voltage-clamp recordings collected from VMHvl^Esr1^ neurons from socially naive, AGG and NON mice (n=11-14 VMHvl^Esr1^ neuron recording per group, collected from 8-10 mice per group, Kolmogorov-Smirnov test, *P =* 0.2780 between socially naive and AGG mice, *P* < 0.0001 between socially naive and NON mice). Right – comparison of sIPSC amplitude in voltage-clamp recordings collected from VMHvl^Esr1^ neurons from socially naive, AGG and NON mice (n=11-14 VMHvl^Esr1^ neuron recording per group, collected from 8-10 mice per group, Kruskal-Wallis one-way ANOVA with uncorrected Dunn’s post hoc test, *P* = 0.8995 between socially naive and AGG mice, *P* = 0.0476 between socially naive and NON mice). ns; not significant, \**P* < 0.05, \*\*\*\**P* < 0.0001. In box-and-whisker plots, center lines indicate medians, box edges represent the interquartile range, and whiskers extend to the minimal and maximal values.

**Fig. S2.**
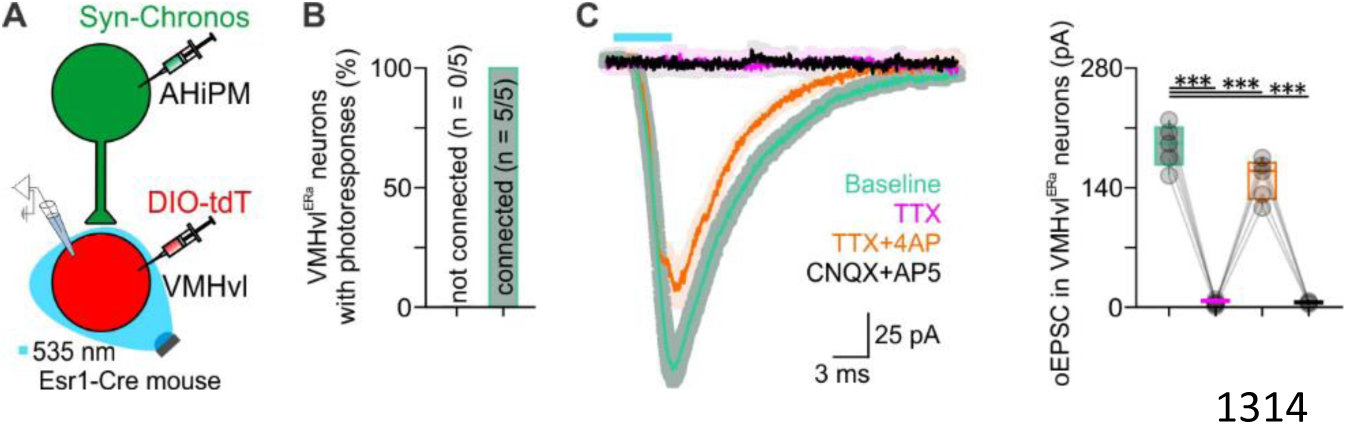
Monosynaptic connectivity between AHiPM and VMHvl^Esr1^ neurons. (A) Schematic illustration of the experimental design, transducing AHiPM neurons with Chronos and optically evoking postsynaptic responses in VMHvl^Esr1^ neurons *ex vivo*. (B) Quantification of VMHvl^Esr1^ neurons with optically-evoked EPSCs (oEPSCs). (C) Averaged amplitudes of oEPSCs evoked on baseline (green), TTX (magenta), TTX + 4AP (orange), and in CNQX and AP5 (black); n=5 brain slices, collected from n=5 mice, one-way ANOVA with Dunnett’s post hoc test, *P* = 0.0002 between baseline and TTX conditions, *P* = 0.0001 between baseline and TTX+4AP conditions, *P* = 0.0002 between baseline and CNQX+AP5 conditions. Shaded region represents the standard error. The vertical scale bar defines current and the horizontal scale bar time. \*\*\**P* < 0.001. In box-and-whisker plots, center lines indicate medians, box edges represent the interquartile range, and whiskers extend to the minimal and maximal values.

**Fig. S3.**
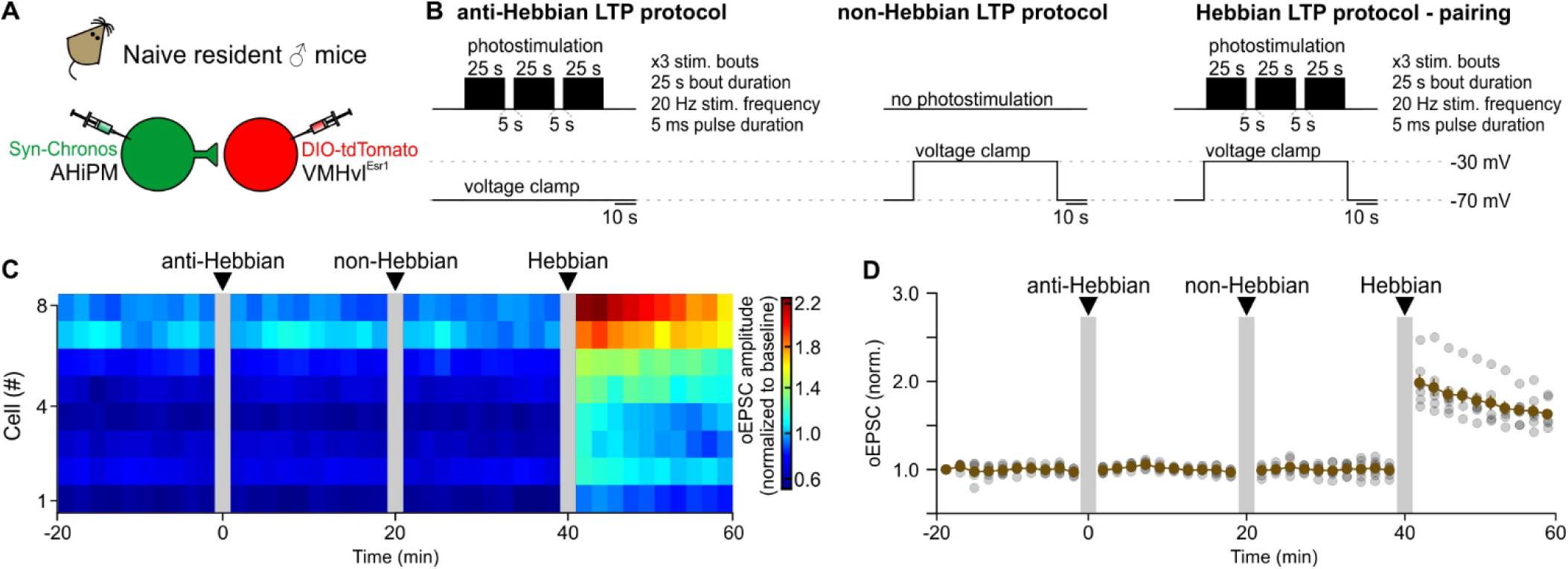
Characterization of LTP-inducing stimulation protocols at the AHiPM➔VMHvl^Esr1^ synapse. (A) Schematic of the experimental design used to identify the appropriate stimulation protocol for LTP induction *ex vivo* in socially naïve mice. (B) Illustration of the experimental protocols tested to to induce LTP in the AHiPM➔VMHvl synapse. (C) Monitoring the optically induced EPSC (oEPSC) prior to, and following application of each of three stimulation protocols (n=8 cells, n=5 socially naïve mice). (D) Alternative quantification/illustration of optically induced EPSC (oEPSC) prior to, and following application of each of three stimulation protocols (n=8 cells, n=5 socially naïve mice – similar to panel C).

**Fig. S4.**
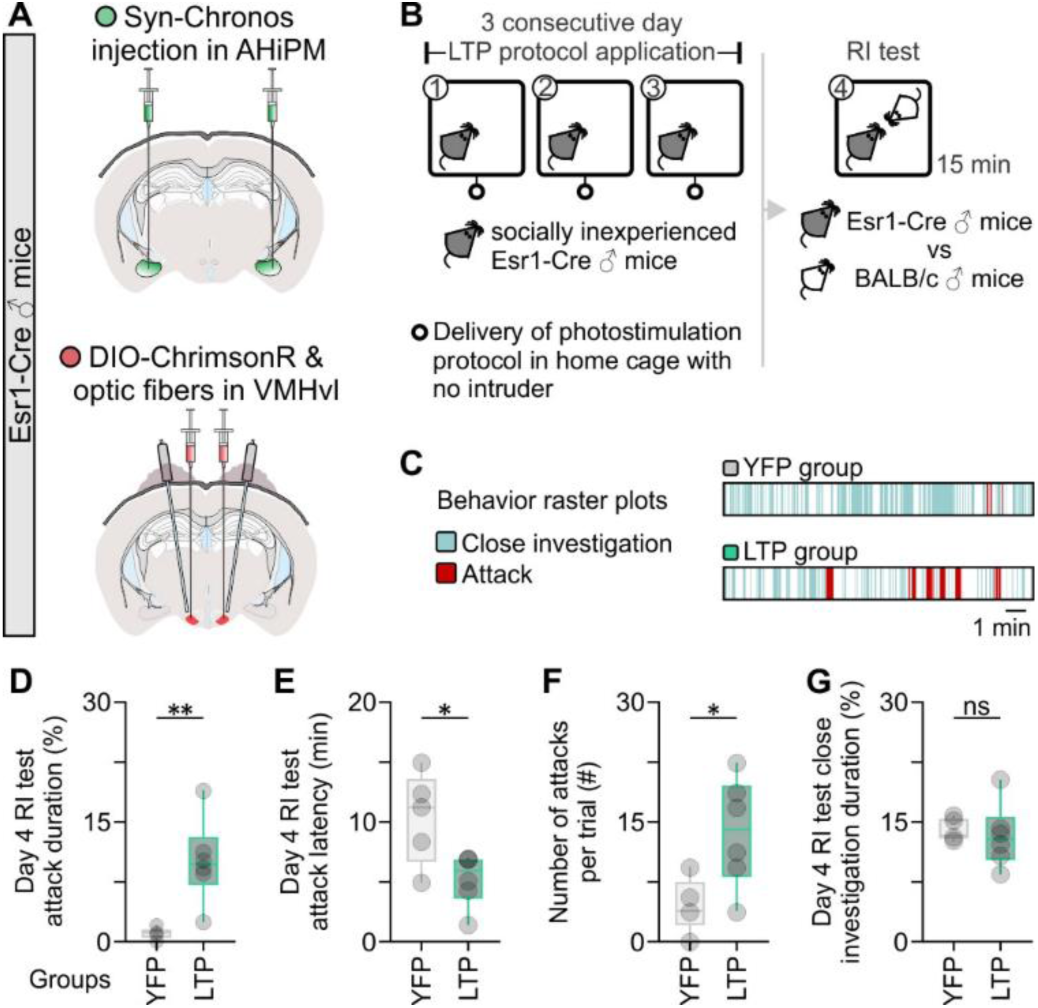
Optogenetic induction of LTP at AHiPM➔VMHvl^Esr1^ synapses in socially naïve mice leads to elevated aggression in the first resident-intruder test. (A) Schematic indicative of the experimental design used to induced hypothalamic LTP in the AHiPM➔VMHvl synapses, via Chronos-eYFP expression in AHiPM, and ChrimsonR expression in VMHvl^Esr1^ neurons. (B) Schematic of the behavior test design used to identify whether induction of LTP in the AHiPM➔VMHvl synapses, influences the innate expression of aggression. (C) Representative behavior raster plots of a control (YFP) and opsin-expressing (LTP) mouse, in the resident-intruder test against a novel BALBc conspecific. (D) Quantification of attack duration (n=5-6 mice per group, two-tailed unpaired *t*-test, *P* = 0.0046 between YFP and LTP groups). (E) Quantification of attack latency (n=5-6 mice per group, two-tailed unpaired *t*-test, *P* = 0.0214 between YFP and LTP groups). (F) Quantification of number of attacks per trial (n=5-6 mice per group, two-tailed unpaired *t*-test, *P* = 0.0235 between YFP and LTP groups). (G) Quantification of close investigation duration (n=5-6 mice per group, two-tailed unpaired *t*-test, *P* = 0.7106 between YFP and LTP groups). \**P* < 0.05, \*\**P* < 0.01. In box-and-whisker plots, center lines indicate medians, box edges represent the interquartile range, and whiskers extend to the minimal and maximal values.

**Fig. S5.**
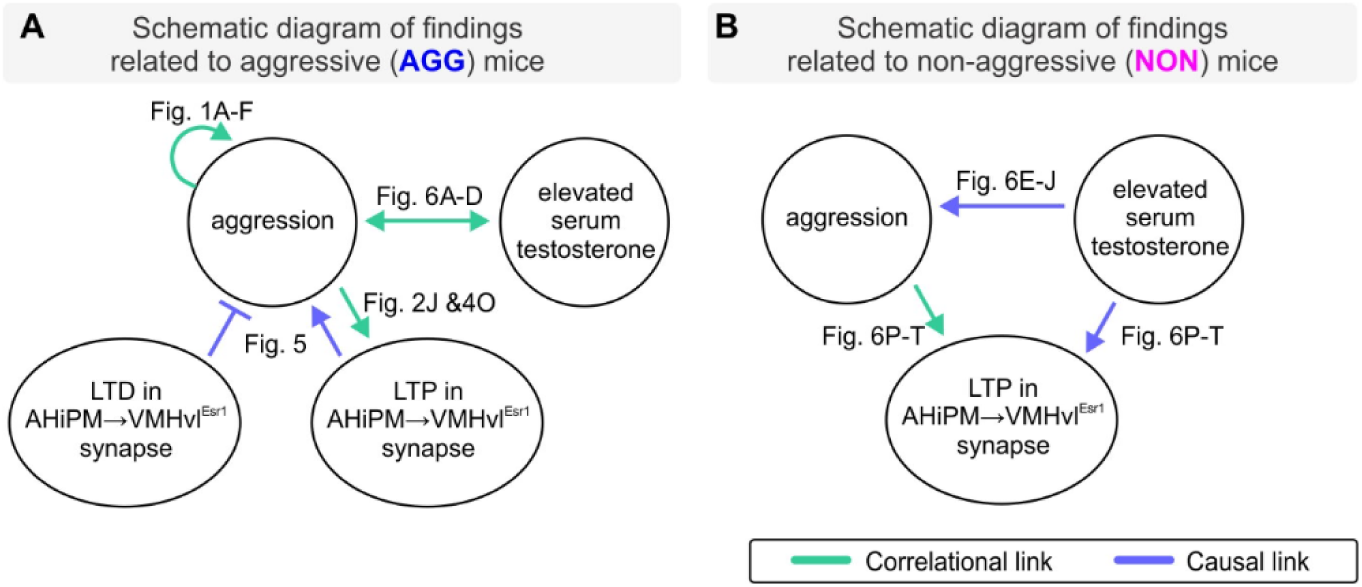
Schematic summary. (A) The schematic summarizes the findings from AGG mice, and the suggested links between aggression, serum testosterone and hypothalamic LTP. (B) Similar to panel (A), but summarizing results from experiments in NON mice - this schematic summarizes the identified links among aggression, serum testosterone and hypothalamic LTP. Our results do not distinguish whether the effect of elevated serum testosterone to increase LTP *in vivo* (Fig. 6P-T) is direct, or rather indirect via an effect to increase aggressive behavior, which in turn enhances LTP. However, exogenous administration of T to NON mice (in the absence of any aggressive experience) enhances LTP amplitude and persistence as tested *ex vivo* (Fig. 6K-O).

## Materials and Methods

**Table S1.**
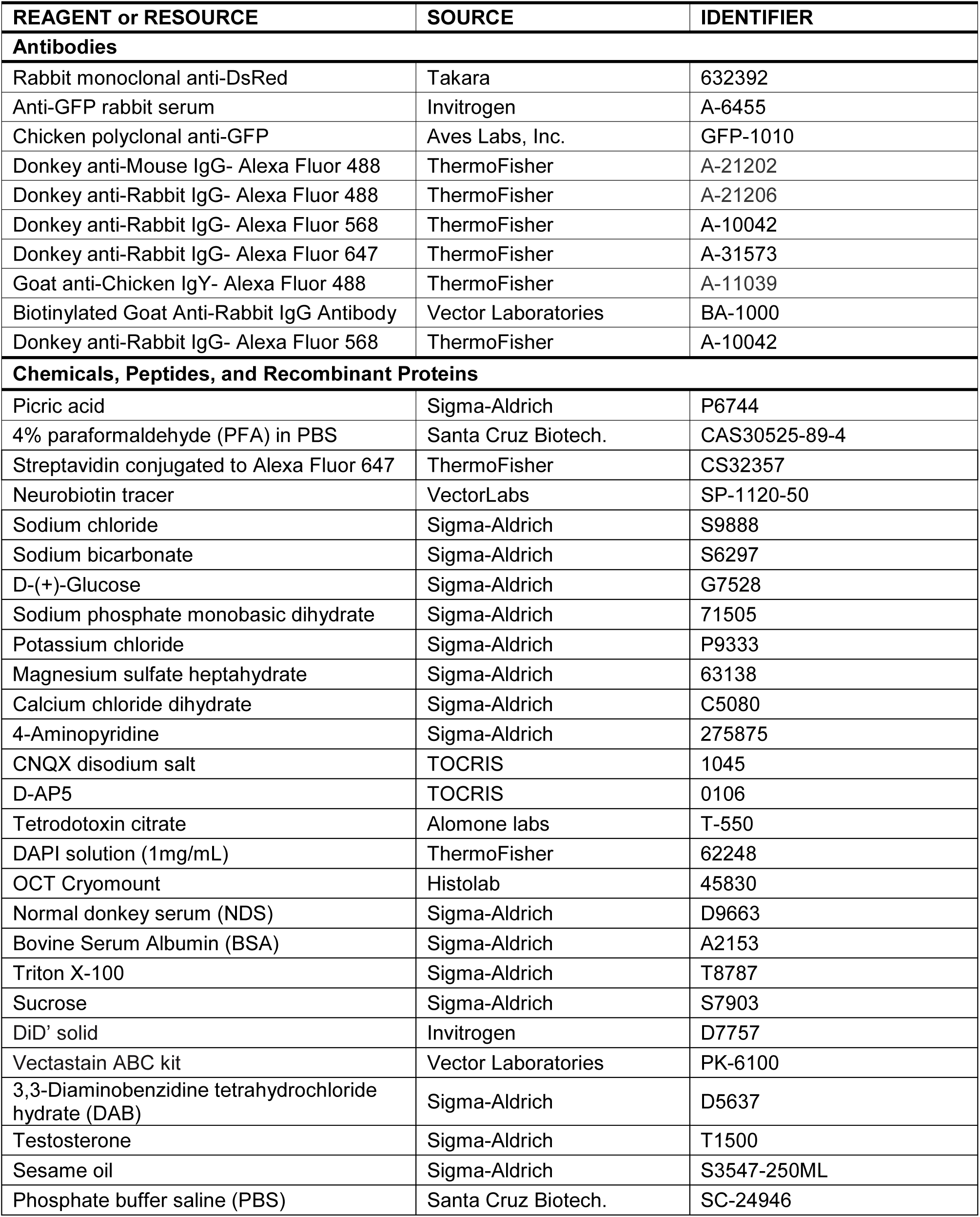
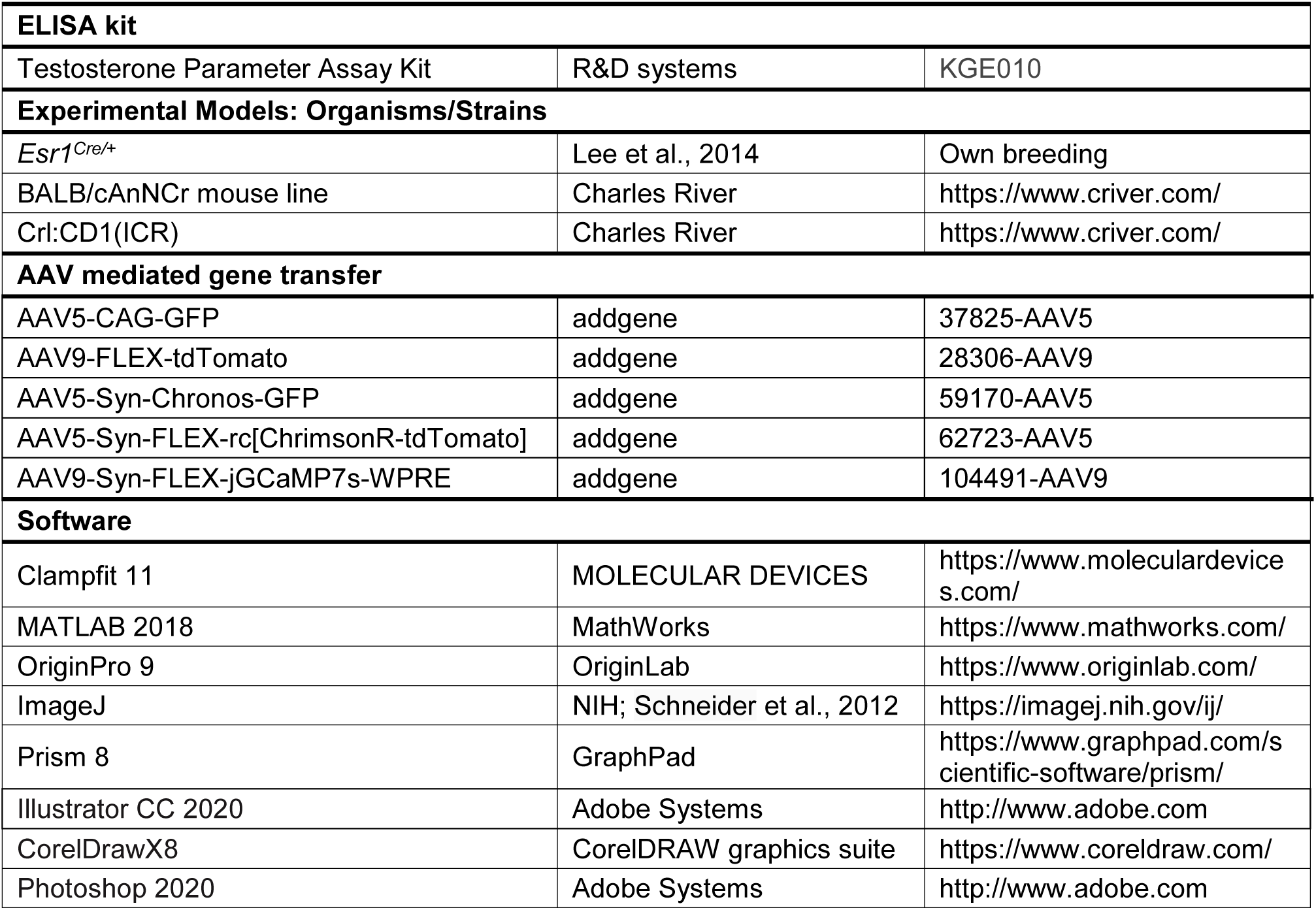
Reagents and resources.

